# Anatomical variability, multi-modal coordinate systems, and precision targeting in the marmoset brain

**DOI:** 10.1101/2022.01.31.478477

**Authors:** Takayuki Ose, Joonas A. Autio, Masahiro Ohno, Stephen Frey, Akiko Uematsu, Akihiro Kawasaki, Chiho Takeda, Yuki Hori, Kantaro Nishigori, Tomokazu Nakako, Chihiro Yokoyama, Hidetaka Nagata, Tetsuo Yamamori, David C. Van Essen, Matthew F. Glasser, Hiroshi Watabe, Takuya Hayashi

## Abstract

Localising accurate brain regions needs careful evaluation in each experimental species due to their individual variability. However, the function and connectivity of brain areas is commonly studied using a single-subject cranial landmark-based stereotactic atlas in animal neuroscience. Here, we address this issue in a small primate, the common marmoset, which is increasingly widely used in systems neuroscience. We developed a non-invasive multi-modal neuroimaging-based targeting pipeline, which accounts for intersubject anatomical variability in cranial and cortical landmarks in marmosets. This methodology allowed creation of multi-modal templates (MarmosetRIKEN20) including head CT and brain MR images, embedded in coordinate systems of anterior and posterior commissures (AC-PC) and CIFTI grayordinates. We found that the horizontal plane of the stereotactic coordinate was significantly rotated in pitch relative to the AC-PC coordinate system (10 degrees, frontal downwards), and had a significant bias and uncertainty due to positioning procedures. We also found that many common cranial and brain landmarks (e.g., bregma, intraparietal sulcus) vary in location across subjects and are substantial relative to average marmoset cortical area dimensions. Combining the neuroimaging-based targeting pipeline with robot-guided surgery enabled proof-of-concept targeting of deep brain structures with an accuracy of 0.2 mm. Altogether, our findings demonstrate substantial intersubject variability in marmoset brain and cranial landmarks, implying that subject-specific neuroimaging-based localization is needed for precision targeting in marmosets. The population-based templates and atlases in grayordinates, created for the first time in marmoset monkeys, should help bridging between macroscale and microscale analyses.

**Highlights:**

- Achieved sub-millimeter localization accuracy of subject-wise brain region
- Propose a dedicated non-invasive multi-modal subject-specific registration pipeline
- Construct brain coordinate system in AC-PC and grayordinate spaces
- Establish multi-modal MRI and CT brain and cortical templates, MarmosetRIKEN20
- Quantify intersubject variabilities in marmoset brain
- Significant bias and uncertainty exist in marmoset stereotactic positioning

## 1. Introduction

Spatial coordinates are a fundamental framework for understanding the brain through mapping cells, architectures and functions. Stereotactic devices are widely used in animal neuroscience and offer a coordinate to map and target specific brain regions (Hardman and Ashwell, 2012; Palazzi and Bordier, 2009; Paxinos et al., 2012; Stephan et al., 1980; Yuasa et al., 2010). Atlases in stereotactic coordinates are commonly based on a single subject’s ex-vivo brain histology data (Bowden and Martin, 2000; Hardman and Ashwell, 2012; Paxinos and Franklin, 2019; Paxinos and Watson, 2017; Saleem and Logothetis, 2006). The assumption behind the stereotactic approach is that each brain structure has consistent coordinates across individuals relative to cranial landmarks (e.g., the bregma, the interaural line, the infra-orbital ridges) (Horsley and Clarke, 1908). While this assumption may hold true in rodents that have low intersubject variability of brain structure and function, it remains unclear in increasingly used small primates such as the New World monkey, common marmoset (*Callithrix jacchus*). For example, the brain volume of marmosets is likely more variable than inbred laboratory strains of rodents: the coefficient of variation (COV) of brain volume is 2.3% in mice (Ma et al., 2008), 3.2% in rats (Hasegawa et al., 2010), and 6.6% in marmosets (Hayashi et al., 2021), and little is known about the positional variability of marmoset cranial and brain landmarks. Intersubject variability of lissencephalic marmoset brains is likely low in terms of neuroanatomy and functional areas but is largely unexplored. It is an intriguing question to ask how brain organisation varies with primate behaviours (Mikula et al., 2007; Pomberger et al., 2019; Yokoyama et al., 2021; De Castro et al., 2021).

The anterior commissure-posterior commissure (AC-PC) coordinate system is another approach originally developed for human neurosurgery and is now routinely used in human neuroimaging. The pioneering work of Tarailach et al. (Talairach et al., 1988) used this approach for deep brain surgery in humans using X-ray ventriculography; brains were standardised to a set of coordinates based in part on the distance between the two landmarks. Reorientation and rescaling of the brain using these intracerebral landmarks was useful to reduce brain variability in size and shape. The origin of AC-PC coordinates is defined in relation to the AC (e.g., its centre or posterior margin) where it intersects the midsagittal plane. The approach was elaborated by improved neuroimaging techniques in particular, magnetic resonance imaging (MRI), which increased the accuracy of brain localization and targeting in both clinical and basic neuroscience. Compensation for subject variability was also elaborated by using automated registration of brain with linear and non-linear algorithm (Evans et al., 1992; Fonov et al., 2011), yielding the Montreal neurological institute (MNI) 152 human template, which is widely used in human neuroimaging. An analogous population-based template and atlas were also developed using MRI in macaques using MRI (Frey et al., 2011; Rohlfing et al., 2012; Seidlitz et al., 2018). Similar approach was also very recently applied for rodents using ex-vivo data at Allen Institute for Brain Science (AIBS) (Hawrylycz et al., 2011; Kuan et al., 2015; Wang et al., 2020).

Neuroimaging-based systems use all structural features (grey matter, white matter, CSF) for registration across subjects but in practice rarely achieve precise alignment of human cerebral cortex owing to the complexity and variability of cortical folding. This has been addressed by accurate cortical segmentation and surface reconstruction by treating the cortex as a 2D sheet-like structure (Dale et al., 1999; Van Essen and Maunsell, 1980). This approach has significantly improved standardisation of cortical anatomy and evaluation of folding pattern and cortical thickness. Glasser et al. (Glasser et al., 2013) further developed a ‘grayordinate’ system which takes into account both the 2D topology of the cortical sheet (ignoring for the moment its finite thickness) and the 3D-volume structure of globular deep brain grey matter structures. A further advance was to apply areal-feature-based alignment using myelin content and fMRI-based resting state networks, which enabled successful definition of cortical areas in in-vivo human brains (Glasser et al., 2016a). Neuroimaging also triggered development of sophisticated targeting systems of brain areas for neurosurgery. However, it has not been established whether a comparably complicated neuroimaging pipeline is needed for a small-brained primate like the marmoset. The stereotactic and AC-PC horizontal planes have been suggested to be parallel to one another in non-human primates (NHPs) (Risser et al., 2019; Saleem and Logothetis, 2006), but this has not, to our knowledge, been critically evaluated. The marmoset’s cortex may be an intermediate between two extremes in mammalian systems neuroscience (i.e., rodents and humans), but it remains unclear which approach is most suitable for achieving maximal accuracy for neuroanatomical and functional targeting.

Here, we explore the variability of cranial landmarks, brain size, and cortical surface landmarks of marmosets to investigate the impact of different coordinate systems and the accuracy of brain localization. Currently available brain coordinate systems in modern neuroscience can be grouped into four types (Table 1): 1) stereotactic coordinates in 3D space mostly based on ex-vivo brain and cranial landmarks in a single-subject (e.g., bregma or interauricular lines) and less commonly in population and/or on the brain landmarks such as AC-PC and midsagittal lines, 2) standard coordinates in a 3D template space based on in-vivo neuroimaging volumes in population mostly oriented using the AC-PC line and midsagittal plane but sometimes using cranial landmarks, 3) coordinates only for the cortical sheet using FreeSurfer based on neuroimaging data, and 4) grey matter-based cortical surface and subcortical volume coordinates (grayordinate) based on neuroimaging data. Rodents, without cortical convolutions, are commonly analysed in stereotactic coordinates created in a single subject (coordinate type 1), whereas recent studies suggest that humans and macaques with cortical folding are most accurately analysed in *grayordinate* system (type 4) (Autio et al., 2020; Donahue et al., 2016; Glasser et al., 2013; Hayashi et al., 2021; Van Essen et al., 2012). We developed a multi-modal marmoset brain targeting system with submillimeter accuracy by utilising a non-invasive head holder, multi-modal marker (Ose et al., 2019), robust image registration initialization using marker-based fiducial registration (MBFR), fine-tuning using a powerful cross-modal registration tool, boundary-based registration (BBR) (Greve and Fischl, 2009), and an analysis pipeline for grayordinates (HCP-NHP pipeline) previously applied in other primate species (human, chimpanzee, macaque) (Autio et al., 2020; Donahue et al., 2016; Glasser et al., 2013; Hayashi et al., 2021). Multi-modal cranial bone, brain template and atlases of the marmoset in the grayordinate system (“MarmosetRIKEN20”) were generated. We also examined the intersubject variability of ‘gold-standard’ stereotactic coordinates and the reproducibility of stereotactic positioning using a stereotactic device in marmosets and evaluated how these are different from AC-PC coordinate system of MarmosetRIKEN20. We demonstrate significant intersubject variability in location of cranial landmarks and cortical surfaces, indicating a need for subject-wise targeting. We also demonstrate a robot-guided neurosurgery with submillimeter accuracy in targeting deep brain structures. We discuss registration accuracy, anatomical variability, templates and atlas, bias between stereotactic and AC-PC coordinates, and neuroimage-based targeting in marmoset.

**Table 1.** Types of brain coordinates used in neuroscience in rodents, non-human primates, and humans.

| Type of brain coordinates | Landmarks | Data used | Subject(s) | References |
| --- | --- | --- | --- | --- |
| 1. Stereotactic | cranial landmarks (bregma, auditory canals, infraorbital line) with an origin of bregma or mid-infraorbital point | Anatomy and histology<br>Ex-vivo MRI | Individual | Rodents (Paxinos and Franklin, 2019; Paxinos and Watson, 2017), Marmoset (Hardman and Ashwell, 2012; Palazzi and Bordier, 2009; Paxinos et al., 2012; Stephan et al., 1980; Yuasa et al., 2010; Woodward et al., 2018), Macaque (Horsley and Clarke, 1908; Paxinos et al., 1999; Saleem and Logothetis, 2006; Hartig et al., 2021; Saleem et al., 2021) |
|  |  | Histology<br>Ex-vivo MRI | Population | Rodents (Kuan et al., 2015; Wang et al., 2020), Marmoset (Majka et al., 2021) |
|  | 3<sup>rd</sup> ventricle, AC-PC and midsagittal interhemispheric plane with an origin of AC | Anatomy and Histology | Individual | Macaque (Francois et al., 1984; Percheron, 1997), Human (Mai et al., 2015; Talairach et al., 1988) |
|  |  | Anatomy and Histology | Population | Rodents (Hawrylycz et al., 2011) |
| 2. Brain volume (3D image-based) | cranial landmarks (bregma, auditory canals, infraorbital line) | Neuroimaging | Individual | Marmoset (Mundinano et al 2016) |
|  |  |  | Population | Marmoset (Hikishima et al., 2011; Risser et al., 2019; Liu et al., 2021) |
|  | AC-PC midsagittal plane | Neuroimaging | Population | Rodents (Papp et al. 2014), Macaque (Frey et al., 2011; Rohlfing et al., 2012; Seidlitz et al., 2018), Human (Holmes et al., 1998; Mazziotta et al., 2001; Rohlfing et al., 2010; Shattuck et al., 2008; Tzourio-Mazoyer et al., 2002) |
| 3. Surface-based | FreeSurfer surface coordinates | Neuroimaging | Population | Macaque (Van Essen and Maunsell 1980), Human (Dale et al., 1999) |
| 4. Grayordinates (3D image and 2D surface-based) | AC-PC, midsagittal plane, grayordinates | Neuroimaging | Population | Marmoset (Ose et al., 2022), Macaque (Autio et al., 2020; Donahue et al., 2016), Human (Glasser et al., 2013) |

## 2. Materials and methods

We used a multi-modal brain targeting system which includes multi-modal markers positioned relative to a head holder, MBFR and BBR for CT and MRI, and a grayordinate analysis pipeline based on high-resolution MRI images. The head holder with multi-modal markers was designed for accurate cross-modal registration between CT and MRI images (Fig. 1). We used a total of 28 common marmosets (*Callithrix jacchus*) (male N = 23, female N = 5, age 5.1 ± 2.7 years and weight 378 ± 59 g; values reported as mean ± SD). For cross-subject standardisation, we generated multi-modal cranial bone, brain and cortical templates of the marmoset (“MarmosetRIKEN20”), which were embedded in AC-PC coordinates and grayordinates using MRI (N = 20) and CT data (N = 10) using the HCP-NHP pipeline. Three animal experiments were conducted: 1) evaluation of the accuracy of the multi-modal brain targeting system and investigation of the intersubject variability of cranial landmarks (N = 10) and cortical landmarks and subcortical structures (N = 20) including cortical sulci (intraparietal (IPS), lateral and calcarine sulci), and subcortical structures, 2) assessment of the reproducibility of stereotactic positioning (N = 5), 3) a proof-of-concept application of the multi-modal brain targeting system to image-guided neurosurgery (N = 1). All the MRI images in this article were obtained using MRI scanner (3-Tesla, MAGNETOME Prisma, Siemens AG, Erlangen, Germany) equipped with a custom-made 16-ch marmoset head coil (Hori et al., 2018). The animals were maintained and handled in accordance with the recommendations of the United States National Institutes of Health. The study was approved by the Institutional Animal Care and Use Committee of the RIKEN Institute in Kobe (MAH28-08).

**Figure 1.**
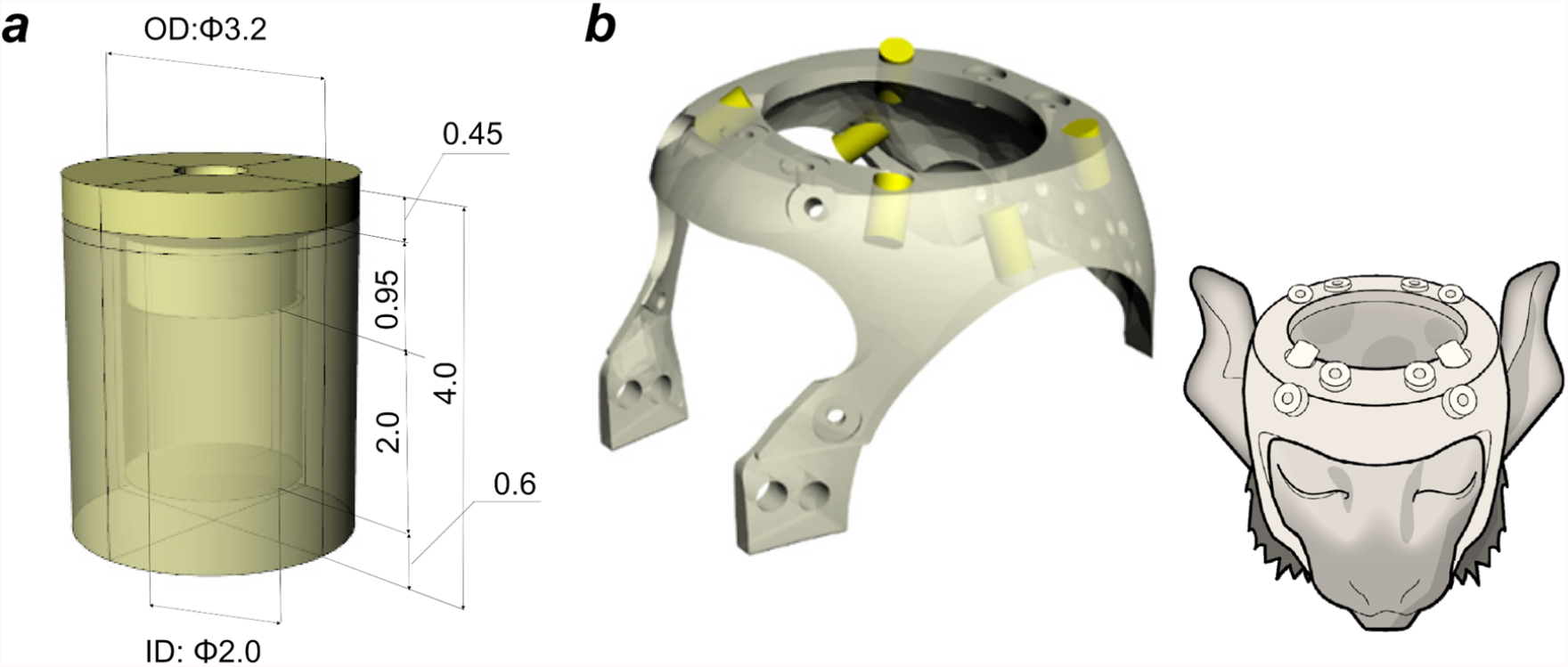
Design and construction of non-invasive head holder equipped with multi-modal markers. **(a)** A multi-modal marker container (outer diameter 3.2 mm, inner diameter 2.0 mm). **(b)** A head holder with six markers (yellow cylinders) (left) was fixed to the marmoset’s head noninvasively (right).

### 2.1 Experiment 1 – registration accuracy of the multi-modal brain targeting system

The registration accuracy of the multi-modal brain targeting system was evaluated using CT and MRI. In each animal, the two scans were performed on the same day on a total of ten marmosets (male N = 7, female N = 3, age 4.0 ± 2.2 years, weight 354 ± 33 g) over a period of ∼2 hours, during which the custom-made plastic head holder with the multi-modal marker was attached to the head. The animals were pretreated with an intramuscular injection of ketamine (30 mg/kg) and dexmedetomidine (5.0 μg/kg) plus atropine sulphate (50 µg/kg). After sedation and respiration were stabilised, the anaesthesia level was maintained with inhaled isoflurane (0.5 %), and intravenous infusion of dexmedetomidine (5.0 μg/kg/hr). Physiology was monitored using a pulse oximeter (7500FO, Nonin, MN, USA); pulse (120 bpm/min) and oxygen saturation (94 ∼ 98 %) were maintained by adjusting the flow rate of anaesthetic gas. Rectal temperature was monitored (Model 1030; SA Instruments, Inc., Stony Brook, NY, USA), and maintained at around 34°C using a custom-made warm water circulation system.

The custom-made plastic head holder with the multi-modal marker consisted of the marker container (Fig. 1a) and head holder (Fig. 1b), which were designed using a 3D software (Rhinoceros 3D v5.0, McNeel & Associates, USA) and produced using a 3D printer (Agillista, Keyence, Osaka, Japan). A compact stereotactic device that couples to the head holder and scanner gantry that we use was also produced (Supplementary Fig. S1). The cylindrical container had outer dimensions of 3.2 mm diameter and 3.55 mm length, inner dimensions of 2.0 mm diameter and 2.0 mm length, and the cap dimensions were 3.2 mm diameter and 0.45 mm length. The container was filled with Tungsten solution (lithium heteropolytungstate [LST]) (Ose et al., 2019) with density (1.9 g/mL) adjusted to be close to the density of cortical bone (White et al., 1989). To prevent evaporation of the liquid, the container was sealed with a cap using a UV-curable resin (Bondic^®^, Laser Bonding Tech, Inc., Aurora, ON, Canada). However, due to the small size of the marker container and the high viscosity of the marker solution, it was challenging to avoid inclusion of air bubbles that reduced the accuracy with which the marker centroid could be determined. Since MBFR requires a minimum of three non-coplanar markers, we included at least six marker containers to ensure a sufficient number of good markers. The shape of the head holder (Fig. 1b) was based on the MR image of a marmoset (male, age 5.1 years, weight 407 g) that represented one of the largest head sizes in our marmoset population (N = 10). To efficiently place the markers around the brain, marker containers were placed so that any combination of three markers was not coplanar (Fig. 1b, yellow cylinders). The head holder was fixed on the animal’s cranium using 8 to 12 resin screws (RENY Pan head machine screw M2.6, SUNCO Industries co.,ltd, Japan) (length: 2-8 mm) tightened to the skin/cranial bone. Before this procedure, the screw points in the skin were locally anaesthetised (lidocaine, 2%, 0.05 ml). This procedure was relatively noninvasive and the skin and bone were not noticeably damaged after removal of the head holder.

After the head-holder was attached, the subject was transported to the MRI scanner (3-Tesla, MAGNETOME Prisma, Siemens AG, Erlangen, Germany) equipped with a custom-made 16-ch marmoset head coil (Hori et al., 2018). T1-weighted (T1w, MPRAGE sequence, TR = 2200 ms, TE = 2.58 ms, TI = 700 ms, flip angle = 8°, averages = 3, scan time = 17.8 min, isotropic voxel size = 0.36 mm, FOV = 70 × 59 × 46 mm) and T2-weighted (T2w, SPACE sequence, TR = 3000 ms, TE = 558 ms, turbo factor = 160, averages = 1, scan time = 6.2 min, isotropic voxel size = 0.36 mm, FOV = 70 × 59 × 46 mm) images were acquired. Next, the animals were transferred from the MRI to the CT. CT was performed on an animal micro CT scanner (R_mCT2, RIGAKU, Japan). The scan parameters were: X-ray voltage of 90 kVp, tube current of 0.2 mA, FOV = diameter 73 mm × height 57 mm, and acquisition time 2.0 min. The CT data was reconstructed (isotropic voxel size = 0.12 mm, matrix = 512 × 512 × 512) with the Feldkamp cone-beam algorithm (Feldkamp et al., 1984). The CT scanner was calibrated to Hounsfield units so that the output images have a value of −1000 for air and 0 for water. This calibration allows us to obtain CT values of −30 to −70 for fat, 20 to 100 for soft tissue, and >1000 for bone (Lev and Gonzalez, 2002).

### 2.2 Experiment 2 – accuracy and reproducibility of stereotactic positioning

Previous studies have not reported the reproducibility of ‘gold-standard’ stereotactic positioning in marmosets. We carried out a rigorous assessment of the bias and reproducibility of stereotaxic positioning using CT imaging (2 female and 3 male, age 5.7 ± 2.4 years, weight 420 ± 42 g). Marmoset heads were fixed to the stereotactic device (Fig. S1), which was custom-constructed to adapt the small bore and FOV of our animal CT scanner (bore diameter = 19 cm, FOV diameter = 7.3 cm). The design of this device was based on commonly available ones (Hardman and Ashwell, 2012; Palazzi and Bordier, 2009; Stephan et al., 1980; Yuasa et al., 2010) and allows secure fixation of the marmoset cranium with ear bars that were firmly inserted into the external auditory canals, eye bars placed above the orbital bones and tooth bars that pushed the upper jaw up to keep tight against the eye bars. This fixation enabled horizontal zero and anterior-posterior zero planes to be perpendicularly oriented with respect to the stereotactic device (Fig. S1a). The tip of the ear bar was set to 2.4 mm diameter, based on the diameter of the external auditory canal (∼2-3 mm) (Kurihara et al., 2019) (Fig. S1c). The animal’s head was fixed to the stereotactic device with ear bars, mouth/tooth bar, and eye bars by an expert experimenter (A.K.) and then CT scanning of the animal’s head and stereotactic device was performed (without the head holder) (Fig. S1d). Then the animal’s head was removed from the scanner gantry and stereotactic device. We repeated the same procedure (positioning, scanning [a 2-min scan], and removal) five times during one session for each animal (a total session duration ≈1 hour). During these experiments the animals were deeply anaesthetised.

### 2.3. Experiment 3 – image-guided neurosurgery

The multi-modal brain targeting system was applied to image-guided neurosurgery in marmosets, and its spatial accuracy to target deep brain structures was evaluated. Specifically, the aim of the surgery was to insert a guide cannula into the caudate nucleus (Cau) or substantia nigra (SN) to administer α-synuclein to induce Parkinson’s Disease-like symptoms in marmosets (Eslamboli et al., 2007; Shimozawa et al., 2017).

Pre-surgery, MRI and CT experiments were performed to guide surgical planning: MRI was used to identify target areas (Cau and SN) and to determine the projection trajectory for surgical operation (Fig. 10a) and reconstruction of cranial surfaces. CT imaging was used to reconstruct the cranial surface and identify initialization landmarks. During MRI and CT scanning, the subject was attached to the non-invasive head holder and registration between images was performed with multi-modal markers as described above.

For surgery, the animal was first anaesthetised with a combination of 0.12 mg/kg medetomidine (0.12 mg/kg), midazolam (0.6 mg/kg), and butorphanol (0.8 mg/kg) (i.m.). The anaesthesia level was maintained with a half volume of the initial anaesthesia dose every two hours. The head of the animal was fixed in the stereotactic cassette of the microsurgical neuronavigation robot (Brainsight, Rogue Research Inc., Montreal, Canada). The robotic components were calibrated in a registration step beforehand (e.g., between two cameras, followed by between the cameras and surgical robot arm). After incision of the skin over the cranium, two cranial landmarks were identified both in a real space with the robot’s laser pointer and in the CT image and used for initialising the registration between the surgical field and the CT image. Then a cranial surface was reconstructed from a point cloud dataset that was scanned using the robot’s laser pointer. Using the CT, the cranial bone was segmented and used to create the CT image-derived cranial surface. The two surfaces were co-registered to each other using the robot’s computer. The target points were defined in the individual brain’s real physical AC-PC native coordinates (SN: X, Y, Z = 2.2, −5.6, −4.9 mm, Cau: X, Y, Z = 3.2, −0.3, 1.9 mm) and the coordinates were registered to robot’s coordinates. The trajectories of the insertion needle were planned using the robot’s controller to define appropriate entry holes and paths for SN and Cau. The cranium was drilled to make these two entry holes (w/ diameter of 1 mm), placed in the medial frontal and parietal areas, to insert guide cannulas into the Cau and SN, respectively. The planned trajectories avoided passing through the lateral ventricles so as to avoid dislocating or deforming the brain (Starr et al., 2010). The dura was pierced with a 26 ga needle, followed by insertion of the guide cannulas and needles into the brain at a velocity of 0.01 mm/sec. The inserted guide cannulas were immobilised to the cranium using resin and the scalp was sutured. The anaesthesia was reversed with atimepazole (antisedan, 0.35 mg/kg, i.m.). All surgical procedures were performed in a sterilised room using sterilised instruments.

Post-surgery, an MRI experiment was performed to evaluate the position of the cannulae in relation to the target (Cau or SN). Because the head together with the attached cannula was larger than the inner size of the marmoset head coil, the scan was performed using a larger 24-channel head coil originally designed for macaque monkeys (Autio et al., 2020). Scanning parameters for the acquired T1w MPRAGE were TR = 2200 ms, TE = 2.23 ms, TI = 700 ms, flip angle = 8°, averages = 6, scan time = 35.0 min, isotropic voxel size = 0.50 mm, and FOV = 56 × 101 × 97 mm. During MRI, anaesthesia was maintained and physiology monitored following the procedures described above (see section 2.1.).

### 2.4. Registration workflow between coordinate systems

As described in the introduction, our primary aim was to establish a multi-modal brain targeting system to enable precise registration between coordinate systems (Fig. 2). The CT images were preprocessed to precisely register to MRI images using MBFR and BBR (Fig 2a, left column). The MR images were processed using the HCP-NHP pipelines (Fig. 2a, right column) (Donahue et al., 2016, Hayashi et al., 2021) (https://github.com/Washington-University/NHPPipelines), which includes three structural pipelines (PreFreeSurfer, FreeSurfer, and PostFreeSurfer) to generate cortical surfaces models of each marmoset hemisphere and register them to a standard grayordinates meshes in CIFTI (Connectivity Informatics Technology Initiative) format (Fig. 2d), a data file format recently standardised to make it easier to work with brain data from multiple disjoint grey matter structures at the same time only including cerebral cortices and other grey matter structures and excluding those not of interest (medial wall, white matter, cerebrospinal fluid) (Glasser et al., 2013).

**Figure 2.**
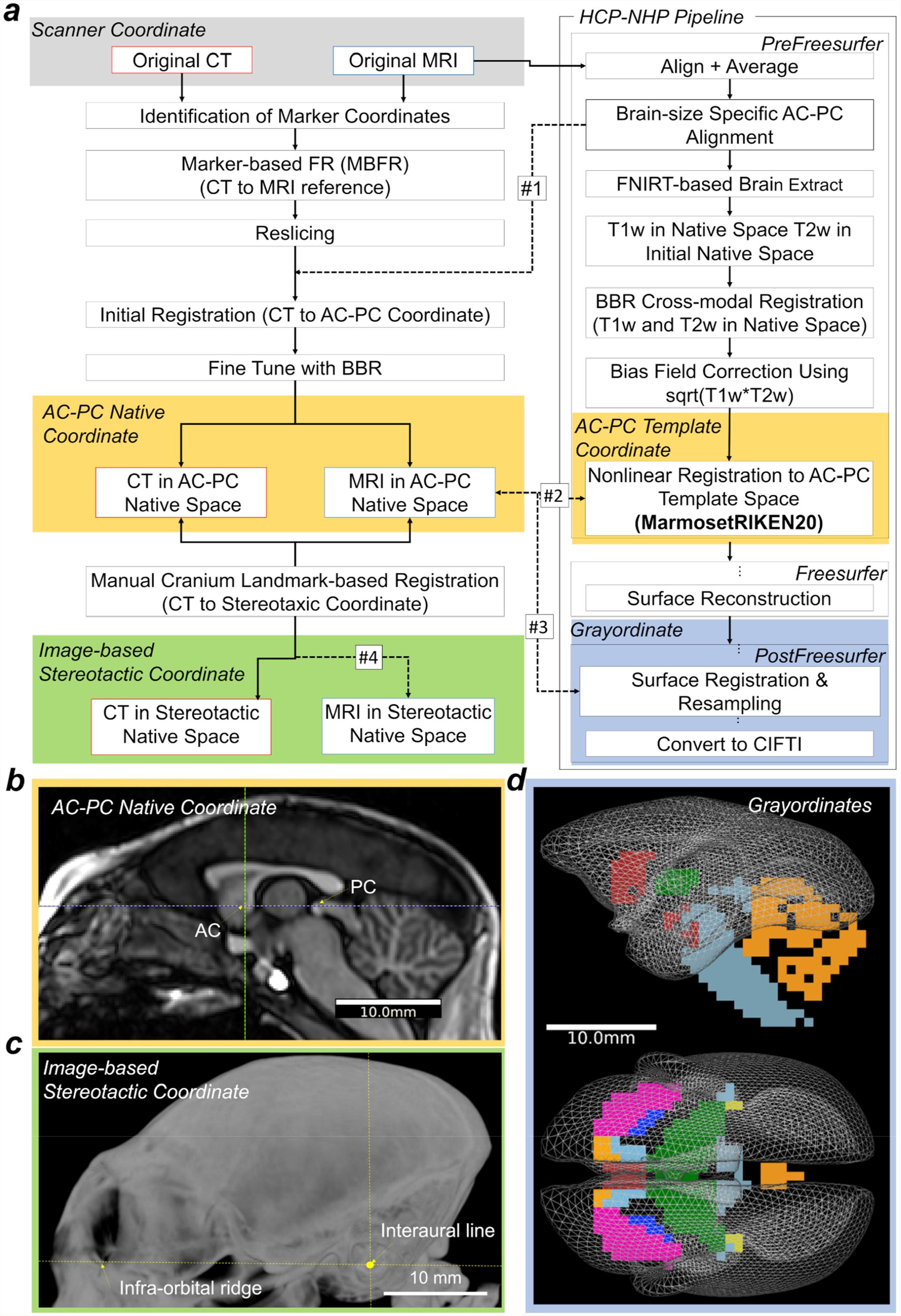
Registration pipeline between coordinates of scanner, anterior-posterior commissure (AC-PC), stereotactic, and grayordinates. **(a)** The workflow describes registration of CT and MR images between different coordinate systems. It contains three main processes: 1) alignment between subject’s CT and MR images, 2) cortical surface reconstruction and registration to AC-PC coordinates using MRI images and the HCP-NHP pipelines, 3) alignment to stereotactic coordinates using the CT image-derived cranial landmarks. From left top: the original CT image was first registered to the original MRI image using a marker-based fiducial registration (MBFR). Then, the CT was transformed to the AC-PC native coordinates and to the AC-PC template coordinates (#1 and #2). On the top right, the original MR image was registered to the AC-PC coordinates in native and the template coordinates using the HCP-NHP pipeline, which generates three transformations: rigid-body matrix (dashed arrows #1), nonlinear warpfield (#2), and resampling to the 4k vertices of the standard-mesh marmoset cortical surface plus the subcortical parcellation (19 parcels, 0.8 mm isovoxels). (#3). The matrix #1 is a linear (rigid-body) registration between the scanner and AC-PC coordinates, the warpfield #2 is a non-linear registration between the AC-PC native and the template coordinates. Finally, the AC-PC-aligned CT in native coordinates was aligned to the image-based stereotactic coordinates using cranial landmarks including external auditory canals and infra-orbital ridge (#4). **(b)** A midline sagittal T1w image aligned in the AC-PC native coordinates. **(c)** Maximum intensity projection of the CT image aligned in stereotactic coordinates, which are orthogonal to the horizontal zero plane passing through both sides of infra-orbital ridges and the interaural line. **(d)** The midthickness surface vertices of 4k CIFTI ‘grayordinates’ in the AC-PC template coordinates. Abbreviations: HCP-NHP, non-human primate human connectome project; BBR, boundary-based registration.

The CT image was aligned to the T2w image in MRI scanner coordinates using MBFR, followed by fine tuned registration with BBR. The workflow of MBFR begins by initial alignment of the original CT to the original T2w image using point registrations (Arun et al., 1987; Kobsch, 1976; Ose et al., 2019), followed by fine-tuning using BBR which uses bone/soft-tissue boundary information of CT in the co-registration process to T2w. For MBFR, each marker’s coordinates in each CT and MRI was identified by calculating the marker centroid after thresholding and binarization for classification of marker and background. The marker coordinates for both CT and T2w were used for registration between CT and T2w using a Kabsch algorithm for point-based registration (Kabsch, 1976). The initial transformation matrix from CT to T2w and resliced CT volume (initialised CT) were generated. We used a custom script, ‘point_reg.py’ for running MBFR, which is made publicly available (https://github.com/RIKEN-BCIL/MultimodalRigidTransform). Then, for BBR, the initialised CT image was threshold at a value of −250, so that images included voxels with soft tissues and bones, then segmented using FSL FAST, and the output of the bone segmentation was fed as a boundary prior into the BBR registration to T2w (in AC-PC native coordinates). The default value of the BBR slope (−0.5) was used. For BBR between CT and T2w, we used ‘epi_reg’ in FSL by specifying CT volume as <whole head T1w image> and T2w image as <EPI image> of epi_reg inputs. We used the T2w for registering CT to MRI images, based on a preliminary analysis that showed the T2w to work better than the T1w because T2w has clear contrast both at the inner and outer boundary of cranial bone. For evaluation of registration, the minimum cost of FSL BBR was calculated using a schedule file ($FSLDIR/etc/flirtsch/measurecost1.sch). The ‘ground truth’ stereotactic coordinates were determined based on the CT image-derived cranial landmarks in each animal (image-based stereotactic coordinates, Fig. 2c), in which the horizontal zero plane passes through both sides of the infra-orbital ridge and the interaural line (the centres of the external auditory canal), and the origin is the point where the interaural line intersects the midsagittal plane (Fig. 2c). To evaluate rotation and translation bias between stereotactic and AC-PC coordinates (Fig. 2b,c), the CT image was also manually aligned using FSL Nudge to ‘ground truth’ image-based stereotactic coordinates using cranial landmarks (i.e., the external auditory canal and infra-orbital margin) and resliced in the stereotactic coordinates in dimensions of X, Y, Z = 254, 254, 136, isotropic voxel size of 0.2 mm, and origin at X, Y, Z = 25.4, −15.4, −4.4 mm (process #4 in Fig. 2b). These ‘ground truth’ image-based stereotactic coordinates were also used as a reference coordinate for measuring the systematic bias and the reproducibility of manual stereotactic positioning (see 2.7).

In the initial step of MRI preprocessing, PreFreeSurfer pipeline, the original T1w and T2w images (in MRI scanner coordinates) were aligned to the individual’s AC-PC native coordinates (Fig. 2b) using a rigid-body transformation (degrees of freedom = 6) with FLIRT in FSL (FMRIB’s Linear Image Registration Tool). The AC-PC line was defined as a line connecting the centre positions of the AC and PC (Schaltenbrand et al., 1977). A CT image in AC-PC native coordinates was generated by applying the transformation matrix (converting original T2w to AC-PC coordinates, Fig. 2, dashed arrow #1) to CT volume realigned to the original T2w by MBFR and BBR (see above) and resampling with spline interpolation. The PreFreeSurfer pipeline calculated the non-linear registration from the T1w in the AC-PC native coordinates to the AC-PC MarmosetRIKEN20 template coordinates described below (Fig. 2, dashed arrow #2) using FNIRT in FSL (http://fsl.fmrib.ox.ac.uk/fsl/fslwiki/FNIRT) and species-specific configuration that scaled size-dependent variables (Hayashi et al., 2021). The linear transformation of original to AC-PC native was combined with a warpfield from AC-PC native to the template to form a single warpfield from the original scanner coordinates to the template, which was then applied to the original CT image with spline interpolation to generate a CT image in the AC-PC template coordinates.

The MarmosetRIKEN20 template was created from 20 marmoset (20 male, age 5.5 ± 2.8 years, weight 380.0 ± 61.0 g) scans using high-resolution T1w images and non-linear registration between subjects using FNIRT. The registrations were iterated three times, with ‘de-drifting’ after each iteration, i.e., removal of the ‘drift’ in mean spatial locations that can occur with registration (Glasser et al., 2016b) in order to approximate the original locations of the marmoset brain. The resultant T1w template volume was embedded in the AC-PC coordinates with dimensions of X, Y, Z = 254, 254, 136, isotropic voxel size of 0.2 mm and origin at X, Y, Z = 25.4, −26.4, −14.8 mm.

The FreeSurfer pipeline used FreeSurfer ver. 5.3-HCP and the T1w and T2w volumes aligned in AC-PC native coordinates for brain signal homogeneity correction, segmentation of white and grey matter and reconstruction of the cortical white and pial surfaces. For this stage, the marmoset brain was scaled five times larger than its original size, so that FreeSurfer could perform high resolution estimation of the subcortical segmentation and then the white matter surfaces. The images were corrected for signal inhomogeneity using a script *IntensityCor*.*sh* using *fsl_anat* in FSL and FreeSurfer normalisation algorithms, *mri_normlize*. Subcortical segmentation was done using a customised probabilistic template of marmosets using FreeSurfer Gaussian Classifier Atlas (Fischl et al., 2002). The white matter segmentation was further tuned using customised white matter skeletons (Autio et al., 2020; Hayashi et al., 2021) that significantly improved the white surface estimation, particularly in the thin white matter blades in the anterior temporal and occipital lobes. The white surface was estimated using a FreeSurfer program customised for the HCP (*mris_make_surface*, in FreeSurfer 5.3-HCP) (Glasser et al., 2013) and then registered across subjects using the FreeSurfer *mris_register* using the marmoset specific option of distance (= 20, default is 5) and maximum search angle (= 50, default is 68) to adjust for the lissencephalic marmoset brain (Hayashi et al., 2021). As a reference for surface registration, a custom population average surface curvature map was created for marmosets using the *mris_make_template* (Fischl et al., 1999). Then, the brain volumes and surfaces were rescaled back to the AC-PC native coordinates, and the pial surfaces were estimated using high-resolution T1w and T2w volumes. We used *mris_make_surface* with the optional argument of max cortical thickness = 3 mm. During pial surface estimation, the corpus callosum was labelled as an area with absent pial surface to avoid the false sulci formation in the retrospleneal region. The PostFreeSurfer pipeline converted the FreeSurfer-based native surface meshes to GIFTI format and resampled them to 164k, 32k, 10k, and 4k meshes in the GIFTI format. The FreeSurfer-based anatomical surfaces (pial and white) were non-linearly warped in 3D to the standard AC-PC template coordinates. The subcortical segmentations of 19 parcels were resampled to a volume with a spatial resolution of 0.8 mm isotropic. The initial cortical surface registration of FreeSurfer concatenated with a group registration across left and right hemispheres (Van Essen et al., 2012) using a multi-modal Surface Matching (MSM) and a folding map (i.e., FreeSurfer ‘sulc’) (Robinson et al., 2014; 2018). The individual to group average registration was performed using a gentle nonlinear registration (*MSMSulc*) based on folding maps (FreeSurfer ‘sulc’) to an average folding map created from 20 marmoset brains. The T1w divided by T2w image was used for generating cortical myelin maps, after removal of any low spatial frequency intensity biases using the smoothed (sigma=3mm) difference with a reference myelin map (Glasser et al., 2013; Glasser and Van Essen, 2011). All the surface metrics (myelin, thickness, sulc, curvature) were resampled into meshes of 164k, 32k, 10k, and 4k surfaces aligned by *MSMSulc* surface registration. The 4k meshes of the left and right hemispheres were combined to make CIFTI grayordiantes consisting of 2840 vertices in each hemisphere (excluding medial wall surface) and 4056 voxels in the subcortical structures (Fig. 2d). The mean spacing of vertices in the 4k mesh was 0.62 ± 0.18 mm on the averaged midthickness surface of the AC-PC template coordinates; median cortical thickness was 1.6 mm (max = 2.6 mm, min = 0.5 mm), and average cortical surface area was 9.9 ± 0.5 cm^2^ (Hayashi et al., 2021).

### 2.5. Analysis of accuracy of the multi-modal brain targeting system

The CT and MR image alignment accuracy was calculated using marker registration error (MRE), which is determined as the root-mean-square of distances (*d*) between the centroid of each corresponding marker point (*1,2*,,…*n*) in the registered CT vs reference MR images.

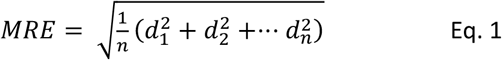

Although there are known limitations in the accuracy of MRE estimation (Danilchenko and Fitzpatrick, 2011; Fitzpatrick, 2009), in our experience it provides a valuable index for the registration accuracy and comparison across registration methods (Ose et al., 2019). The MRE was compared between MBFR, normalised mutual information (NMI), BBR and a combination of MBRF and BBR. The registration was performed using either of two ways to specify the region of interest: full FOV and a partial FOV thresholded to remove background (partial FOV and threshold). Thresholds of 2 and −1000 were used in MRI and CT, respectively, to set background (air) to zero. The partial FOV tightly enclosed the marmoset head.

Registration was reported as ‘failed’ when the MRE was greater than 1 mm, since errors this size are unacceptable for many purposes. The probability of registration failure was calculated by dividing the number of failed trials by the total number of registrations. The failure probability was evaluated and compared across different registration methods.

### 2.6. Analysis of intersubject variability of brain and cranial landmarks

To evaluate intersubject variability of the cranial volume and shape, we investigated cranial contours in both AC-PC native and AC-PC template coordinates. We also evaluated the pitch rotation angle (frontal downward direction) of the AC-PC coordinates compared with the ‘true’ (image-based) stereotactic coordinates. The brain and intracranial volume were obtained in AC-PC native coordinates from segmented T1w and CT images, respectively. In addition, stereotactic surgery reference (bregma) and cephalometric (inion, rhinion, and zygion) points were identified from the CT in the native AC-PC coordinates, based on a previously described method (Paxinos and Franklin, 2019). The landmark variations were displayed with respect to the average cortical white and pial surfaces generated from a large population of marmosets (N = 20). We also calculated the maximum intensity projection of the CT images so that we could visualise where the bregma is located and define the x-y plane of each subject.

We also investigated cortical surface landmark intersubject variability. Specifically, we focused on the intraparietal sulcus (IPS) because it is a recognizable landmark in marmosets (Fig. 7b) (Chaplin et al., 2013; Paxinos et al., 2012). We quantitatively defined the presence of the IPS based on the values of the FreeSurfer ‘sulc’ measure in each subject’s 32k mesh, and identified the local minimum in a region of interest (IPS ROI), which comprises four intraparietal areas (anterior intraparietal [AIP], medial intraparietal [MIP], lateral intraparietal [LIP], and ventral intraparietal [VIP] areas). These intraparietal regions were created from a volume representation of the Paxinos atlas which was non-linearly warped to the AC-PC template coordinates and mapped onto the surface (Paxinos et al., 2012). The IPS was considered to be present if the minimum of sulc was lower than −0.37 in the ROI. The 3D coordinates of the vertex with the minimum sulc was identified on the midthickness surface in the subject’s 32k native (AC-PC) coordinates. We also identified the coordinates of the calcarine and lateral sulcal terminations (extrema) in the T1w AC-PC native coordinates in all the animals (N = 20), and calculated average and standard deviations.

We also evaluated the average and variability for the volumes of subcortical regions across subjects. The high-resolution non-linear registration warp field was applied to the 11 subcortical atlas regions (amygdala, habenular nucleus, inferior colliculus, internal pallidum, lateral geniculate nucleus, medial geniculate nucleus, nucleus accumbens, red nucleus, stria terminalis, subthalamic nucleus) and embedded in the T1w AC-PC native coordinates of each animal. The volume and the coordinate of the centroid was evaluated in each animal and the mean and standard deviation were calculated across subjects (N = 20). Six distances between the landmarks including brain length, brain width, anterior-posterior length of corpus callosum, anterior tip of frontal pole to anterior tip of temporal pole, anterior tip of temporal pole to posterior tip of lateral sulcus, anterior to posterior tip of calcarine sulcus (see Supplementary Fig. S2) were also evaluated across subjects.

### 2.7. Analysis of precision and reproducibility of stereotactic positioning

The reproducibility of manual positioning of the cranium within the stereotactic device (device-based stereotactic coordinates) was investigated using repeated mounting (N = 5, n = 5; total 25 experiments) and bias was determined by comparison with the ‘true’ (image-based) stereotactic coordinates. After each mounting, the marmoset head and stereotactic device was scanned using CT. All the CT images were registered using a rigid-body transformation by weighting the mask for the stereotactic device, resulting in the same location relative to the stereotactic device, which we refer to as device-based stereotactic coordinates that include experimenter’s fixation errors. Then, the error between device-based and ‘true’ (image-based) stereotactic coordinates were estimated by using FSL Nudge and a rigid-body transformation (a rotation and translation for each of three axes, see Fig. 4a). The mean and 95% confidence interval of transformation parameters were calculated using averaged data of repeated positioning as a representative value for each animal (N = 5) and analysed to assess the bias of device-based stereotactic coordinates by using a Wilcoxson signed rank test. To evaluate reproducibility of the device-based stereotactic coordinates, the intraclass correlation coefficient (ICC, type 1,1) (Shrout and Fleiss, 1979) of repeated measures of transformation parameters was calculated using the R package “psych” (William, 2020).

### 2.8. Analysis of accuracy of neurosurgical localization

The operational accuracy (to insert a guide tube into the brain) was estimated by the target error defined as the distance between the pre-operational plan in the SN and the postoperative trajectory of the guide cannula. The target location was determined using the preoperative MR image, and the trajectory of the guide cannula was evaluated using the postoperative MR image. The orthogonal distance between the preoperative target point and actual operative trajectory was calculated to estimate the target error.

## 3. Results

### 3.1. Registration accuracy and precision between multi-modal images

The CT to MRI MRE (see Eq. 1) was compared among registration methods (NMI, BBR and MBFR) as shown in Figure 3. The MRE was very small using MBFR w/o BBR (0.15 ± 0.04), whereas those of software-based registrations (NMI or BBR) were significantly larger using all FOV setups (*p*-values ≤ 0.05, one-way repeated measures ANOVA) (Fig. 3a). The large MRE errors (> 1.0 mm) can be ascribed to initialization failure (Greve and Fischl, 2009). Initialization failure probability (with a threshold at MRE > 1.0 mm) was very high for software-based methods, ranging from 0.3 to 1.0 (0.7 in NMI and full FOV; 1.0 in NMI and partial FOV; 0.3 in BBR and full FOV; 0.7 in BBR and partial FOV), whereas it was zero using MBFR. The high initialization failure of software-based methods may be ascribed to the sphere-like shape of the marmoset head. The failed registration of BBR resulted in higher values of the minimum cost (> 0.8), whereas those of the successful registrations were reasonably small (mean 0.56 ± 0.06 in full FOV; 0.59 ± 0.06 in partial FOV). When the BBR followed the MBFR, there were no failures in any subjects, and it resulted in a small number of min costs (0.68 ± 0.05). Importantly, while this MBFR+BBR approach resulted in relatively higher values of MRE than with a MBFR only approach, this is likely due to circularity in defining the MRE based on the MBFR landmarks, as the registration of the brain and cranium were improved for MBFR+BBR compared with MBFR by visual inspection of all ten subjects (Fig. 3b,c). Therefore, these findings demonstrate the robustness and accuracy of the MBFR+BBR approach as compared with software-only registrations.

**Figure 3.**
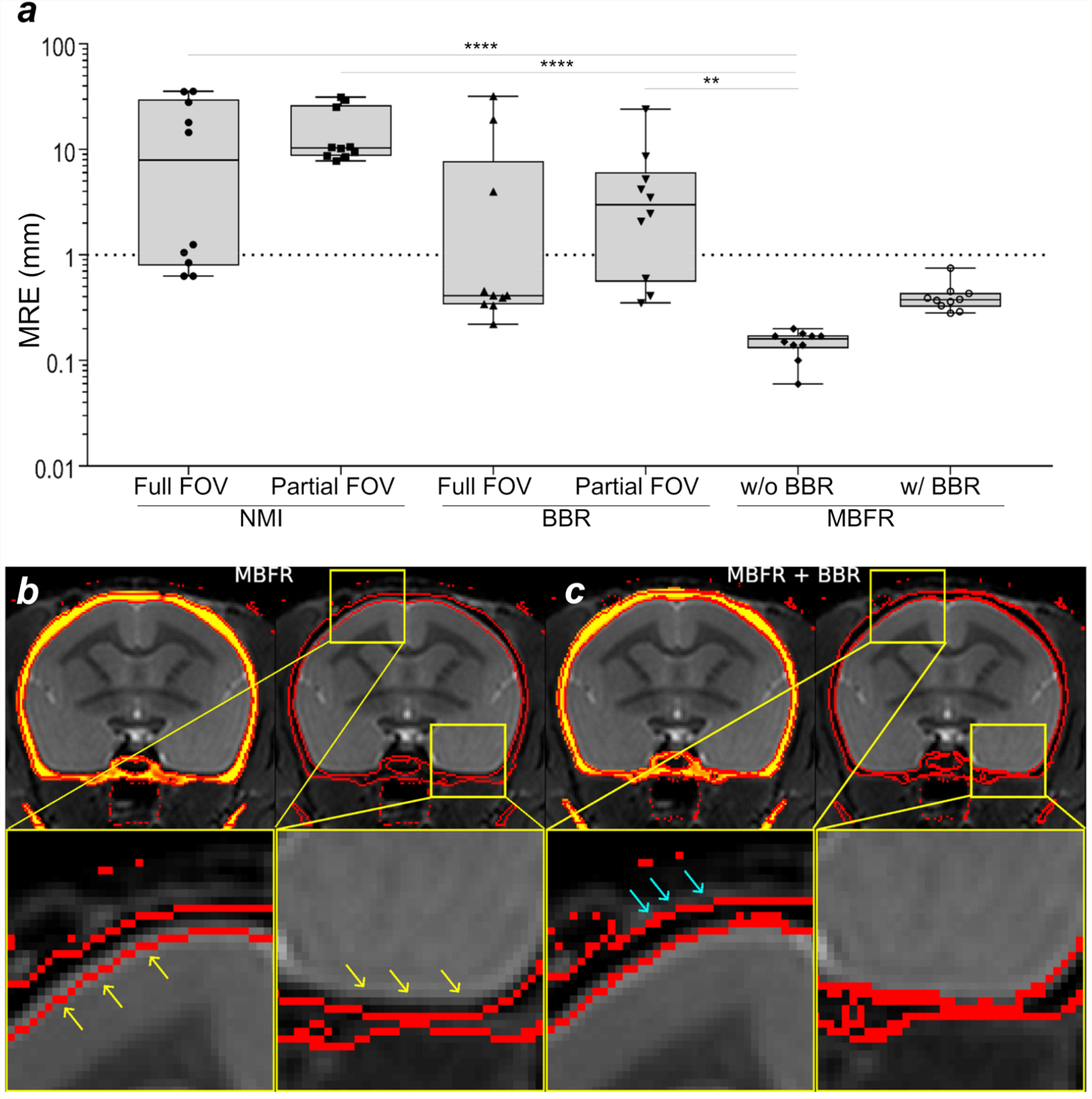
Comparison of accuracy and precision between registration methods. **(a)** Comparison of the marker registration error (MRE) between normalized mutual information (NMI), boundary-based registration (BBR) marker-based fiducial registration (MBFR) and MBFR followed by BBR methods. Registration between CT and MR images of marmosets (N = 10) was performed with NMI and BBR using full and partial field-of-view (FOV), and MBFR. Note that MBFR provided significantly smaller MRE in comparison to other approaches (*p*-values ** ≤ 0.01, **** ≤ 0.001). The threshold of failure was defined as 1 mm (dotted line). **(b)** The results of MBFR between CT and T2w. Note the small but clearly visible error in registration (yellow arrows) of inner cranium boundary (red line) extends into the brain parenchyma, **(c)** MBFR followed by BBR tuneup shows more precise alignment of cranium inner surface to the outline of cortical surface. Note also that the outer cranium boundary was also well aligned to the signal loss boundary of the T2w (aqua arrows). Study ID: (CT: 19042302, MRI: A19042302).

Although the MBFR showed 100% success rate of the registration and alignment was fairly good, a careful inspection revealed subtle (potential) misalignments at the cranium-brain interface in some of the subjects (Fig. 3b, see magnified snippets). This is probably because accuracy of the marker-based registration should depend on the accurate identification of the marker centroid (see section 4.1). To overcome this limitation, we applied a second stage of registration using BBR which registers cranium boundary and head image and found excellent image alignment for brain and cranium boundaries (Fig. 3c). Thus, two-stage registration using MBFR followed by BBR achieved the most robust and accurate registrations between CT and MRI images.

### 3.2. Intersubject variability of cranial contours and coordinate bias between stereotactic and AC-PC space

The measured volume was 6,180 ± 524 mm^3^ and 6,912 ± 470 mm^3^ for brain and cranial cavity, respectively. The intersubject variability (coefficient of variation) was 8.5% and 6.8% respectively. Cranium contours were also highly variable across subjects both in ‘true’ image-based stereotactic coordinates (Fig. 4a) and AC-PC native coordinates (Fig. 4b). Notably, the cranial positions in the image-based stereotactic coordinates (Fig. 4a) are significantly rotated from the AC-PC native coordinates (Fig. 4b), with frontal regions downwards at an average pitch (rotation around X-axis) of 10.0 ± 1.3° (N = 10, p = 0.02). Rotation bias was also found in the roll (Y-rotation) albeit by a much smaller angle (0.6 ± 0.1°, p = 0.002) (see mid panel for coronal section in Fig. 4a), whereas bias in yaw (Z-rotation) was negligible (0.2 ± 0.6°, p = 0.32) (see right panel for axial sections in Fig. 4a). After non-linear registration across subjects, cranial contours were reasonably well registered across subjects in the AC-PC template coordinates (Fig. 4c). There seems to be asymmetry of auditory canals ‘with respect to’ the symmetric brain, as was shown by non-zero roll (0.6 ± 0.1°) and yaw (0.2 ± 0.6°) between “true” stereotactic coordinates (i.e., symmetrical ear canal) and AC-PC template (i.e., symmetrical brain), which is likely due to asymmetry of the auditory bone canals relative to the brain.

**Figure 4.**
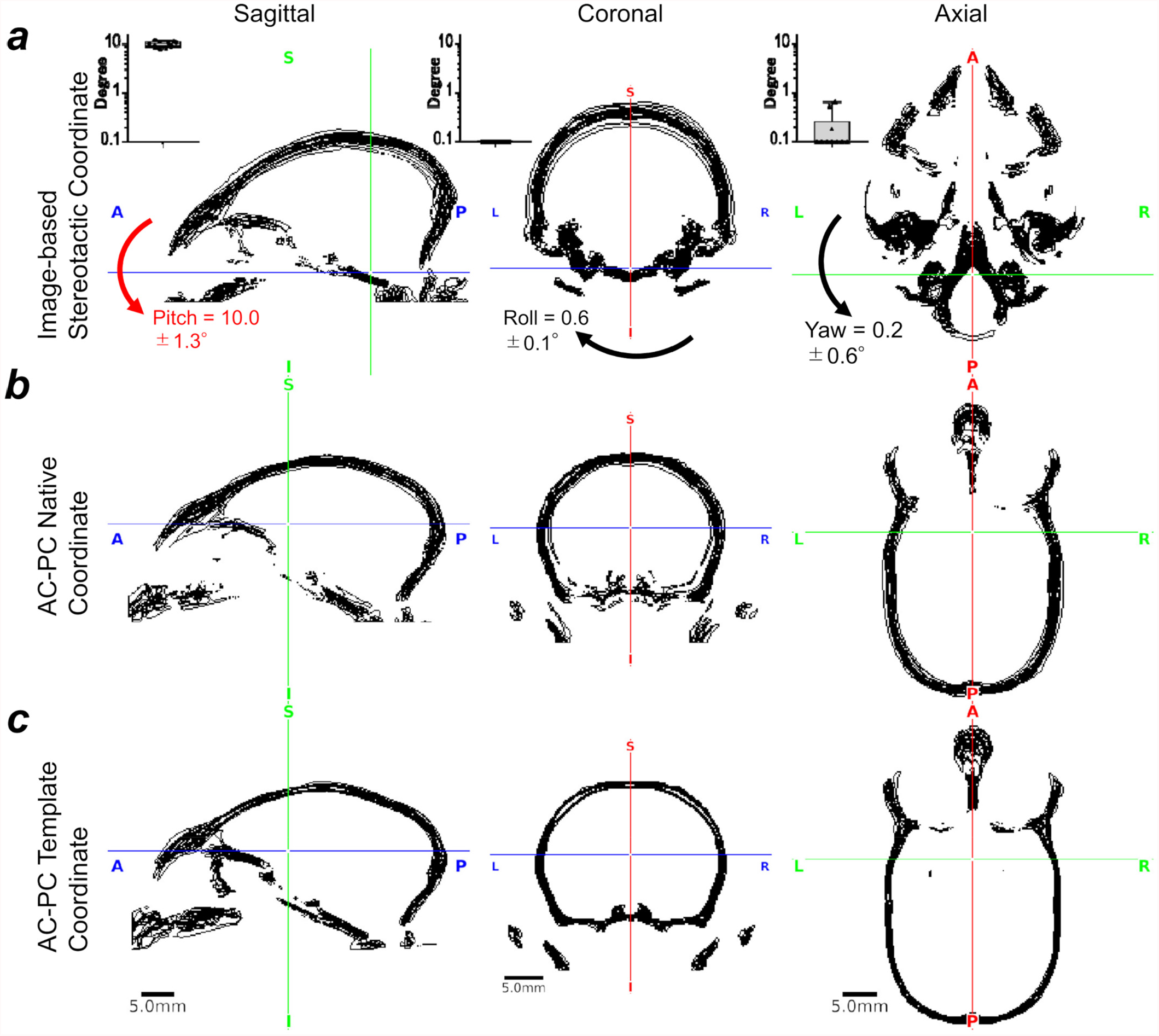
Marmoset intersubject variability in AC-PC and stereotactic coordinates. The cranial bone contours, obtained using CT (N = 10), are displayed in black lines **(a)** in the ‘true’ (image-based) stereotactic coordinates, **(b)** the AC-PC native coordinates and **(c)** in the AC-PC template coordinates. The crosshair shows the origins (mid interauricular line in stereotactic and the centre of AC in AC-PC coordinates). Cranial position was tilted downwards (pitch) from AC-PC native coordinates to stereotactic coordinates (10.0 ± 1.3°, p < 0.0001), whereas roll, 0.6 ± 0.1° and yaw 0.2 ± 0.6°. Note that the cross-subject variability of the cranial contours in stereotactic coordinates are relatively small in the areas close to the origin but large in the distant areas from the origin, particularly in the dorsal convexity of the cranium. In contrast, the extent of the cross-subject variability is similar in the dorsal, ventral and fronto-occipital areas in AC-PC coordinates. The cross-subject variability in the AC-PC template was smaller than the AC-PC native coordinates, suggesting successful non-linear registration of the brain and cranium between subjects. Study ID: (CT: 19021301, 19022601, 19022602, 19022603, 19022701, 19022702, 19042301, 19042302, 19060401, 19060402)

### 3.3. Stereotactic positioning bias and reproducibility

The errors of manual positioning of the cranium are shown in Figure 5. When evaluated in the manually positioned device-based stereotactic coordinates (Fig. 5a), the variability of the cranial contours is caused by both intrasubject (e.g., experimenter’s positioning reproducibility) and intersubject variability (e.g., animal’s cranial shape and size). Therefore, the variability of the contours is not only found in the dorsal convexity of the cranium but also in the areas around the auricular canals (Fig. 5a). In contrast, the contour errors in the image-based stereotactic coordinates almost exclusively demonstrate intersubject variability and no intrasubject variability can be seen (Fig. 5b). Figure 5c shows the bias of the manually positioned device-based stereotactic coordinates with respect to the ‘true’ image-based coordinates. There were trends of rotation biases in pitch (1.6 ± 0.4°, p = 0.06) and roll (rotation in Y-axis) (1.1 ± 0.5°, p = 0.06), but not yaw (rotation in Z-axis) (−0.2 ± 0.7°, p = 0.8) (Fig. 5c, left panel). These biases may be coming from the instability of the fixation device at the skin and soft tissue in the orbital ridge bone and the auditory canal bone. Translation exhibited no significant bias (Fig. 5c, right panel). The ICC (1,1), a measure of the intrasubject reproducibility, was poor in X-rotation (−0.11) and Z-translation (0.33), moderate in X and Y translations (0.80 and 0.84, respectively) and excellent in Y and Z rotations (0.93 and 0.98, respectively) (Fig. 5e). The poor reproducibility in x-rotation and z-translation is likely due to the imperfect accuracy in positioning the fixation device at the orbital ridges or to variability of the skin and soft tissue.

**Figure 5.**
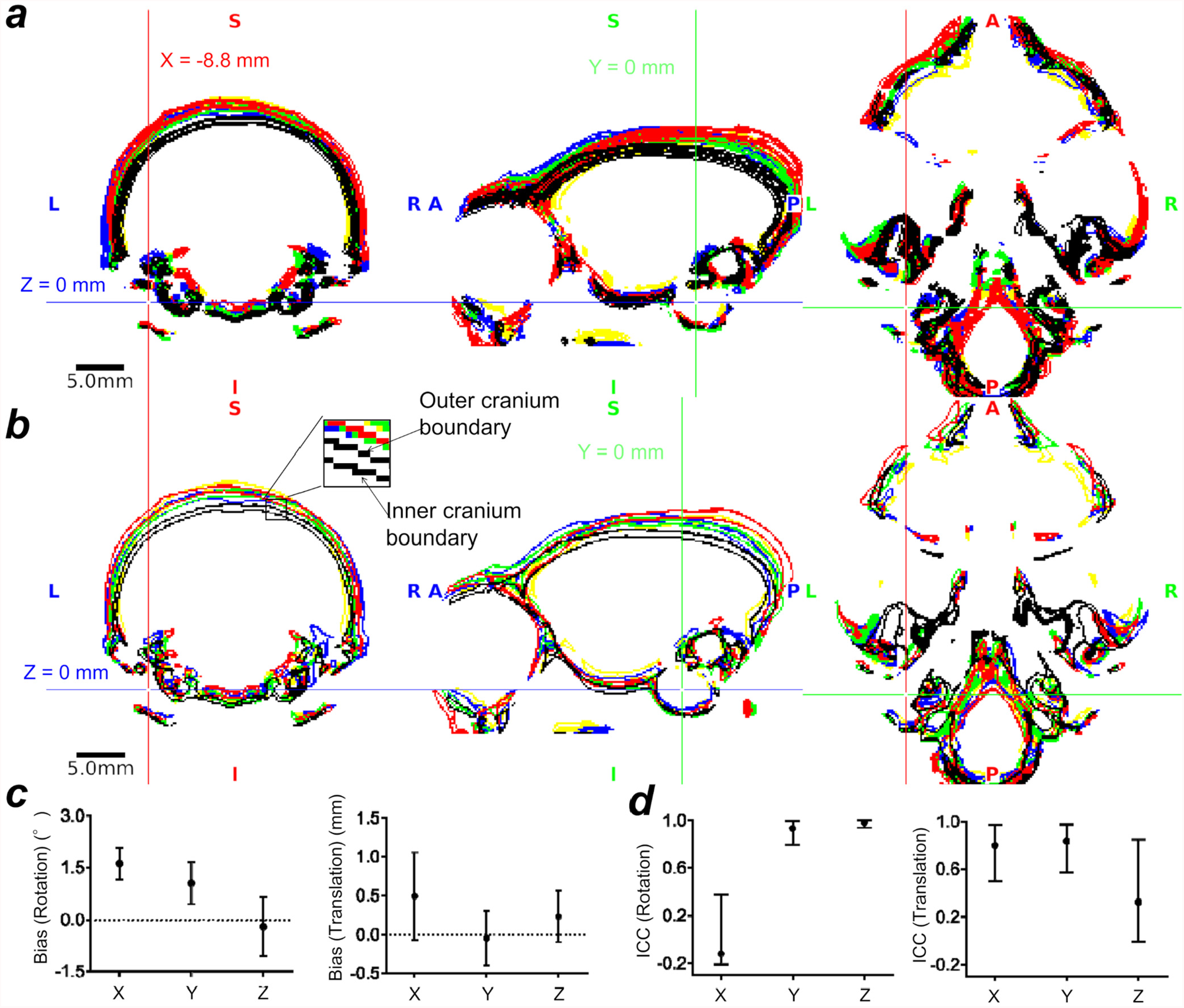
The stereotactic positioning bias and reproducibility. **(a)** Cranial contours of five repeated positioning in five subjects in manually positioned device-based stereotactic coordinates. Each colour indicates a subject’s cranial contour, and the crosshair is placed at the centre of the tip of the ear bar. The cranial contours demonstrated variability both in intrasubject (e.g., experimenter’s positioning reproducibility) and in intersubject (e.g., animal’s cranial shape and size). Note that the variability of the contours is particularly evident in the dorsal convexity of the cranium, which is distant from the ear canal. **(b)** Cranial contours of five subjects in ‘true’ image-based stereotactic coordinates. Note that the locations of the cranial contours are highly variable across subjects although the fixation points (ear canal and orbital ridge) are well colocalized across subjects. In **(a)** and **(b)**, the colours indicate different subjects. Inset denotes two lines in each colour, one for the outer cranium and the other for inner cranium boundary. **(c)** Bias of the device-based stereotactic coordinates in rotations (left) and translations (right) relative to the ‘true’ image-based stereotactic coordinates (N = 5). **(d)** Intra-class correlation (ICC(1,1)) of rotations and translations of device-based stereotactic coordinates (N = 5, n = 5, total 25 experiments).The error bars indicate the 95% confidence interval. Study ID: (CT: 19051001, 19082102, 19082103, 19082104, 19082105)

### 3.4. Intersubject variability of landmark locations, distances, and volumes of interest in AC-PC native coordinates

Intersubject variations of the positions of the bregma in top-to-bottom view of cranium are shown in Fig. 6 and other cranial and cortical landmarks in Fig. 7a, respectively (N = 10). Among the investigated landmarks, the largest variation was unexpectedly found in the bregma in the anterior-posterior direction (±1.0 mm, SD, Table 2, Fig. 6,7) in AC-PC coordinates. This size of the variability is notable as it represents over 5% of the average marmoset brain length in AP-direction (31 ± 0.8 mm, N = 10). Also we note that the shape of the bregma was highly variable across subjects (see Fig. 6 for maximum intensity projections of the cranium for all the subjects examined). In addition, moderately high variability (±1.0-1.1 mm) was found in the inion and rhinion in the superior-inferior direction (Z) and the right zygion in the anterior-posterior direction (Y).

**Table 2.** Intersubject variability of cranial and cortical landmark coordinates (N = 20)

|  | Side | AC-PC Native Coordinates |  |  | Stereotactic Coordinates |  |  |
| --- | --- | --- | --- | --- | --- | --- | --- |
|  |  | X | Y | Z | X | Y | Z |
| Cranial |  |  |  |  |  |  |  |
| Bregma | - | -0.1 ± 0.1 | -7.0 ± 1.0 | 10.0 ± 0.3 | -0.3 ± 0.2 | 6.0 ± 1.1 | 20.4 ± 0.5 |
| Inion | - | 0.1 ± 0.1 | -23.0 ± 0.5 | -3.0 ± 1.1 | -0.2 ± 0.2 | -12.0 ± 0.7 | 10.5 ± 1.2 |
| Rhinion | - | -0.1 ± 0.3 | 19.6 ± 0.9 | -3.2 ± 1.0 | 0.1 ± 0.2 | 29.9 ± 0.8 | 2.9 ± 0.5 |
| Zygion | L | -15.1 ± 0.4 | -0.2 ± 0.8 | -10.0 ± 0.5 | -15.1 ± 0.5 | 9.4 ± 0.6 | -0.4 ± 0.1 |
|  | R | 14.9 ± 0.7 | -0.1 ± 1.1 | -10.0 ± 0.5 | 14.9 ± 0.5 | 9.2 ± 0.8 | -0.3 ± 0.3 |
| Cortical |  |  |  |  |  |  |  |
| IPS | L | -5.5 ± 0.5 | -10.3 ± 0.8 | 5.5 ± 0.3 | -5.5 ± 0.5 | 2.3 ± 0.8 | 16.4 ± 0.3 |
|  | R | 4.8 ± 0.5 | -10.0 ± 0.6 | 5.6 ± 0.4 | 4.8 ± 0.5 | 2.5 ± 0.7 | 16.4 ± 0.3 |
| Posterior tip of lateral sulcus | L | -7.8 ± 0.6 | -6.8 ± 0.9 | 4.3 ± 0.7 | -7.7 ± 0.6 | 5.5 ± 0.8 | 14.7 ± 0.8 |
|  | R | 7.8 ± 0.5 | -6.5 ± 0.7 | 4.1 ± 0.4 | 7.8 ± 0.5 | 5.7 ± 0.7 | 14.4 ± 0.4 |
| Anterior tip of calcarine sulcus | L | -1.8 ± 0.3 | -11.4 ± 0.3 | 1.6 ± 0.7 | -1.8 ± 0.3 | 0.5 ± 0.3 | 12.9 ± 0.4 |
|  | R | 1.8 ± 0.3 | -11.5 ± 0.3 | 1.7 ± 0.2 | 1.8 ± 0.3 | 0.4 ± 0.3 | 12.9 ± 0.2 |
| Posterior tip of calcarine sulcus | L | -2.2 ± 0.3 | -20 ± 0.4 | -1.2 ± 0.4 | -2.2 ± 0.3 | -8.6 ± 0.4 | 11.6 ± 0.4 |
|  | R | 2.0 ± 0.3 | -20.2 ± 0.3 | -1.3 ± 0.5 | 2.0 ± 0.3 | -8.7 ± 0.4 | 11.5 ± 0.4 |

X, Y, and Z indicate the coordinates for cranial (N = 10) and cortical (N = 20) landmarks in the AC-PC native and stereotactic coordinates (mm; mean ± SD). Note that the left intraparietal sulcus (IPS) was identified in seven subjects and the right IPS in three subjects. The coordinates of cranial landmarks were transferred from AC-PC native coordinates to stereotactic coordinates using the subject-specific warp file and the cortical landmarks were transferred using the template warp file.

**Figure 6.**
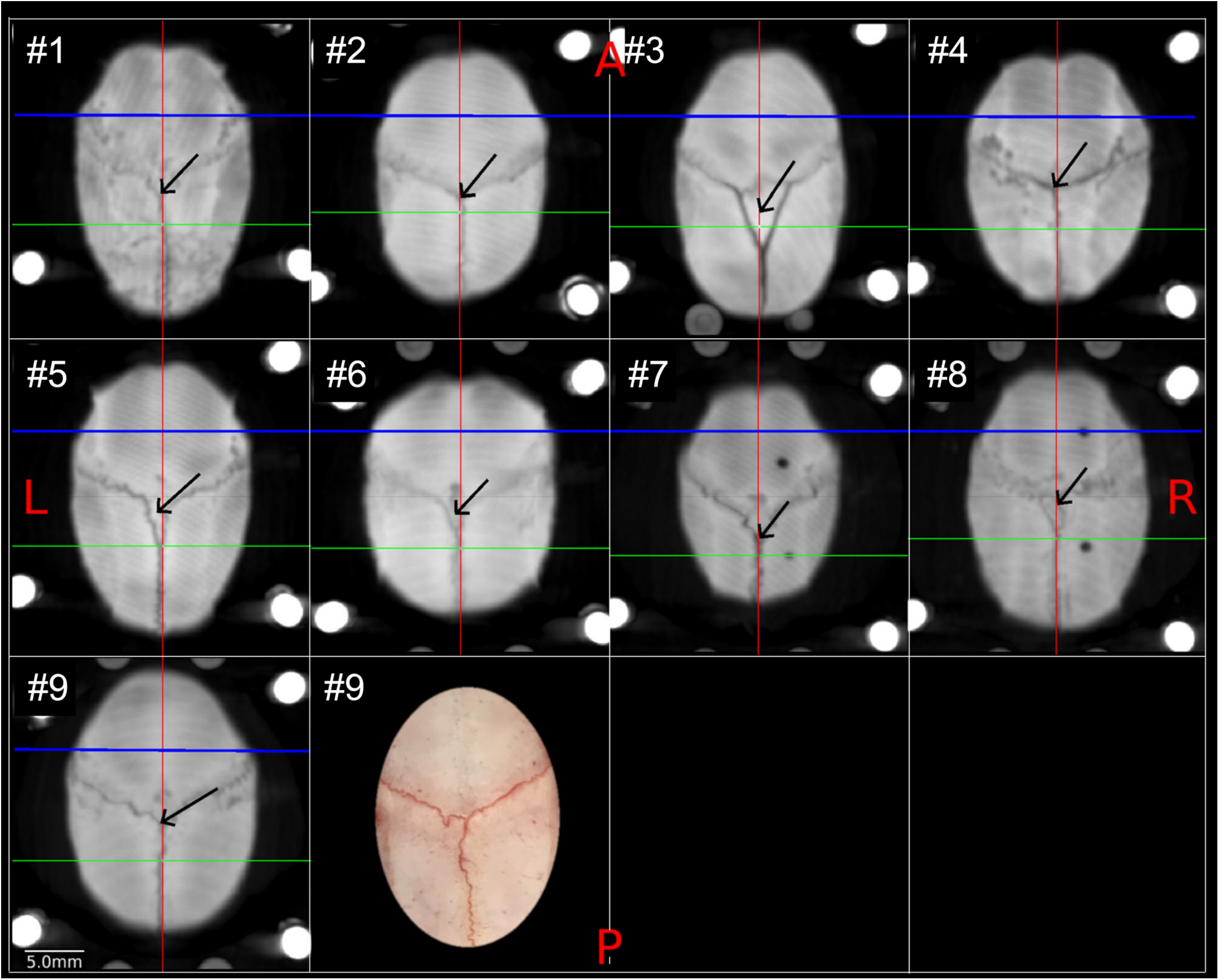
Variability of cranial sutures and bregma in marmoset. Each panel shows the maximum intensity projection of CT images in the x-y plane (N = 9), demonstrating the coronal and sagittal cranial sutures and the estimated location of bregma (black arrow). The lines indicate the y-axis in the midline (red line), x-axis of the interauricular line (green) and AC origin (blue). The bregma was defined as the midpoint of the curve of best fit along the coronal suture (Paxino and Franklin 2019). Note that the coronal suture splits in some of the subjects (e.g., #3) and in other subjects turn sharply (e.g., #5, 6) just before midline, which makes determining the bregma ambiguous based on extrapolation of coronal sutures from lateral to medial. The right panel of the animal #9 shows a photograph of the dorsal view of exposed cranium and sutures. In one of the subjects (#10), cranial sutures and bregma could not be reliably identified (data not shown). Study ID: (CT: 19021301, 19022601, 19022602, 19022603, 19022701, 19022702, 19042301, 19042302, 19060401)

**Figure 7.**
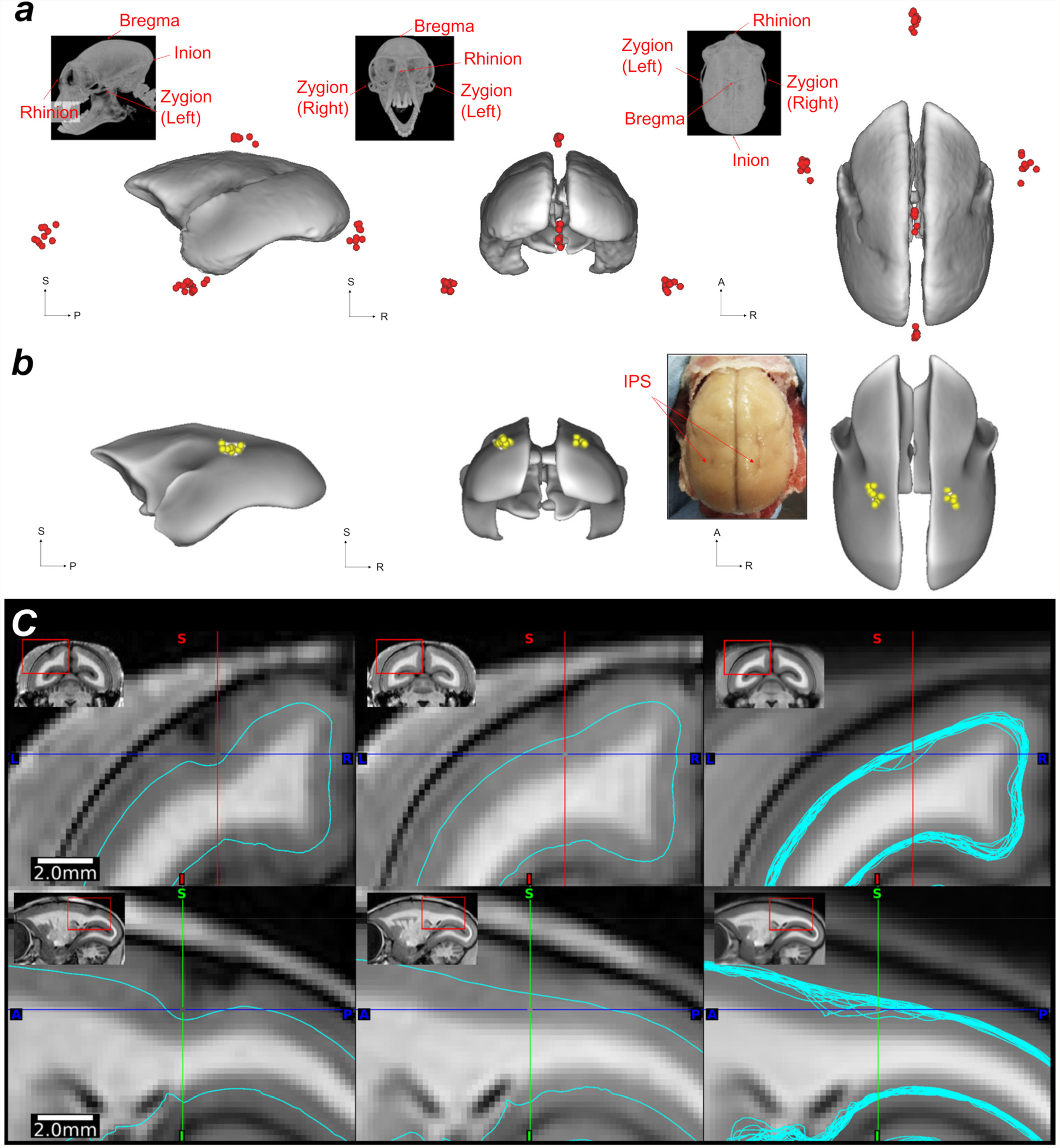
Landmark variability across common marmosets. **(a)** Cranial landmark (bregma, inion, rhinion, zygion-left, and zygion-right.) intersubject variability (N = 10) displayed on the AC-PC template pial surface. **(b)** The deepest points of the intraparietal sulcus (IPS) and their intersubject variability displayed on the white matter surface of the AC-PC template (white nodes, N = 20). The inset shows the macroscopic view of the ex vivo brain, demonstrating the IPS in both hemispheres. **(c)** Midthickness surfaces in the AC-PC native coordinates in an animal with moderate IPS (left) and with negligible IPS (middle). The right panel shows all the midthickness surfaces (N = 20) warped into the AC-PC template coordinates (right), demonstrating the cross-subject variability of midthickness surfaces around IPS even after non-linear volume registration to the AC-PC template coordinates. Study ID: (CT: 19021301, 19022601, 19022602, 19022603, 19022701, 19022702, 19042301, 19042302, 19060401, 19060402, MRI: A19021301, A19022601, A19022602, A19022603, A19022701, A19022702, A19042301, A19042302, A19060401, A19060402, A17051101, A17041201)

Intersubject variation of cortical landmarks (i.e., IPS) is shown in Fig. 7b,c. Among all animals (N = 20), the IPS was not identified in both hemispheres in all the animals: 5% of animals (N = 1) had IPS in both hemispheres, 10% (N = 2) only in the right, 30% (N = 6) only in the left. The locations of IPS deepest points were variable across subjects (Fig. 7b and Table 2, N = 20) variability was moderate in anterior-posterior direction (2SD = 1.2 and 1.6 mm in right and left hemispheres, respectively), followed by left-right (2SD = 1.0 and 1.0 mm), and inferior-superior (0.6 and 0.6 mm, respectively) directions, corresponding to 4 −5%, 4%, and 3% of the average brain lengths in each direction. An example of the variability of IPS is shown in two representative individuals (Fig. 7c). In the animal with a clear IPS (Fig.7c, left panel), the IPS was easily identifiable and the deepest point of cortical midthickness is easily identifiable (blue line), whereas in subject #2 (Fig.7c, middle panel), identification of IPS was challenging, and the cortical midthickness (aqua) was relatively smooth and shallow. Note that the deepest points of the IPS varied by approximately 0.5 mm in the vertical direction when these two subjects’ midthickness surfaces were displayed over the cross-subject average volumes (Fig.7c, right panel) after a warpfield from AC-PC native to the AC-PC template coordinates. We also assessed the location of the end of the lateral fissure, anterior and posterior ends of the calcarine sulcus (Table 2). The variability of y-dimension of lateral fissures were comparable with those of IPS, while variability of the coordinates of calcarine sulcus ends were slightly smaller in both native coordinates of AC-PC and stereotactic spaces (Table 2).

Intersubject average and variations in the volume of brain and subcortical structures are shown in Table 3. The coordinates of the centre of the gravity in AC-PC native coordinates, as well as distances of the brain landmarks, are shown. The results indicate that the overall subject variability in regional volume was 5 to 12% by coefficient of variation (COV), and the variability in distance of several landmarks was 3 to 8%.

**Table 3.** The intersubject variability in volumes of brain and subcortical structures and distances of landmarks of interest (N = 20)

| Volumes / distances of interest | Side | mean ± SD (mm<sup>3</sup> / mm)<br>(COV %) | Coordinates of center-of-gravity |  |  |
| --- | --- | --- | --- | --- | --- |
|  |  |  | X | Y | Z |
| Whole brain | - | 7683.2 ± 440.2 (6) | - | - | - |
| Amygdala | L | 43.2 ± 2.3 (5) | -5.2 ± 0.2 | -0.4 ± 0.4 | -5.2 ± 0.3 |
|  | R | 42.8 ± 2.0 (5) | 5.2 ± 0.2 | -0.4 ± 0.3 | -5.2 ± 0.3 |
| Habenular nucleus | L | 1.5 ± 0.1 (8) | -0.9 ± 0.1 | -6.5 ± 0.3 | 1.2 ± 0.1 |
|  | R | 1.4 ± 0.1 (8) | 0.9 ± 0.1 | -6.5 ± 0.3 | 1.2 ± 0.1 |
| Inferior colliculus | L | 8.3 ± 0.5 (6) | -2.6 ± 0.1 | -10.3 ± 0.3 | -2.7 ± 0.3 |
|  | R | 8.4 ± 0.7 (8) | 2.5 ± 0.1 | -10.2 ± 0.2 | -2.7 ± 0.3 |
| Internal pallidum | L | 5.0 ± 0.4 (8) | -4.2 ± 0.1 | -2.1 ± 0.3 | -1.3 ± 0.2 |
|  | R | 4.3 ± 0.3 (7) | 4.2 ± 0.2 | -2.0 ± 0.3 | -1.4 ± 0.2 |
| Lateral geniculate nucleus | L | 13.9 ± 0.8 (6) | -6.4 ± 0.1 | -5.5 ± 0.3 | -1.6 ± 0.2 |
|  | R | 14.1 ± 1.0 (7) | 6.5 ± 0.2 | -5.5 ± 0.2 | -1.7 ± 0.2 |
| Medial geniculate nucleus | L | 2.6 ± 0.2 (9) | -4.9 ± 0.1 | -6.4 ± 0.3 | -2.1 ± 0.2 |
|  | R | 2.5 ± 0.2 (9) | 4.8 ± 0.2 | -6.4 ± 0.2 | -2.2 ± 0.2 |
| Nucleus accumbens | L | 17.8 ± 1.0 (6) | -2.3 ± 0.1 | 2.2 ± 0.4 | 0.1 ± 0.2 |
|  | R | 16.5 ± 1.2 (7) | 2.3 ± 0.1 | 2.2 ± 0.4 | 0.1 ± 0.2 |
| Red nucleus | L | 1.0 ± 0.1 (12) | -1.2 ± 0.1 | -6.3 ± 0.3 | -4.1 ± 0.3 |
|  | R | 1.0 ± 0.1 (10) | 1.3 ± 0.1 | -6.3 ± 0.3 | -4.1 ± 0.3 |
| Stria terminalis | L | 2.8 ± 0.2 (7) | -1.7 ± 0.1 | -0.8 ± 0.3 | 0.7 ± 0.2 |
|  | R | 2.8 ± 0.3 (11) | 1.6 ± 0.1 | -0.8 ± 0.3 | 0.7 ± 0.1 |
| Subthalamic nucleus | L | 0.7 ± 0.0 (6) | -3.3 ± 0.1 | -3.6 ± 0.3 | -2.2 ± 0.2 |
|  | R | 0.8 ± 0.1 (10) | 3.4 ± 0.1 | -3.6 ± 0.3 | -2.2 ± 0.2 |
| Brain length | - | 32.7 ± 0.8 (3) | - | - | - |
| Brain width | - | 24.2 ± 0.6 (3) | - | - | - |
| Anterior-posterior length of corpus callosum | - | 13.5 ± 0.5 (4) | - | - | - |
| Anterior tip of frontal pole - anterior tip of temporal pole | L | 12.7 ± 0.4 (3) | - | - | - |
|  | R | 12.8 ± 0.5 (4) | - | - | - |
| Anterior tip of temporal pole - posterior tip of lateral sulcus | L | 13.7 ± 1.0 (8) | - | - | - |
|  | R | 13.4 ± 0.6 (5) | - | - | - |
| Anterior to posterior tip of calcarine sulcus | L | 9.2 ± 0.5 (5) | - | - | - |
|  | R | 9.2 ± 0.5 (6) | - | - | - |

### 3.5. Averaged AC-PC template coordinates of marmoset brain and cranium

Using the multi-modal brain targeting system we generated multi-modal MRI (N = 20) and CT (N = 10) AC-PC templates (Fig. 8). These templates provide detailed positional relationships between the brain and cranium. Interestingly, the CT template also reveals physiological calcifications, which were colocalized in the globus pallidus and dentate nuclei bilaterally in the MRI template (Fig. 8a, green and cyan arrow). Figure 8b shows the subcortical parcellations of MarmosetRIKEN20 (version 1.0), which included 21 subcortical grey matter regions (caudate, putamen, external segment of globus pallidus, internal segment of globus pallidus, nucleus accumbens, stria terminalis, claustrum and end-piriformis, thalamus, habenular nucleus, red nucleus, subthalamic nucleus, substantia nigra, superior colliculus, inferior colliculus, lateral geniculate nucleus, medial geniculate nucleus, amygdala, hippocampus, periaqueductal grey, dorsal raphe nucleus, cerebellar cortex) and anterior and posterior commissures. The templates also included T1w-divided-by-T2w myelin map (Fig. 8c) and surface version of the marmoset cortical parcellation atlas of Paxinos, Watson, Petrides, Rosa and Tokuno (Paxinos et al., 2012) including 116 cortical areas (Fig.8d). The distribution of the cortical myelin showed that high myelin signal is colocalized with the parcellations of MT, somatomotor (4ab) and somatosensory (3a, 3b) areas and visual cortex (V1), and modestly high myelin with the frontal eye field (FEF, area 8av) (Fig. 8e).

**Figure 8.**
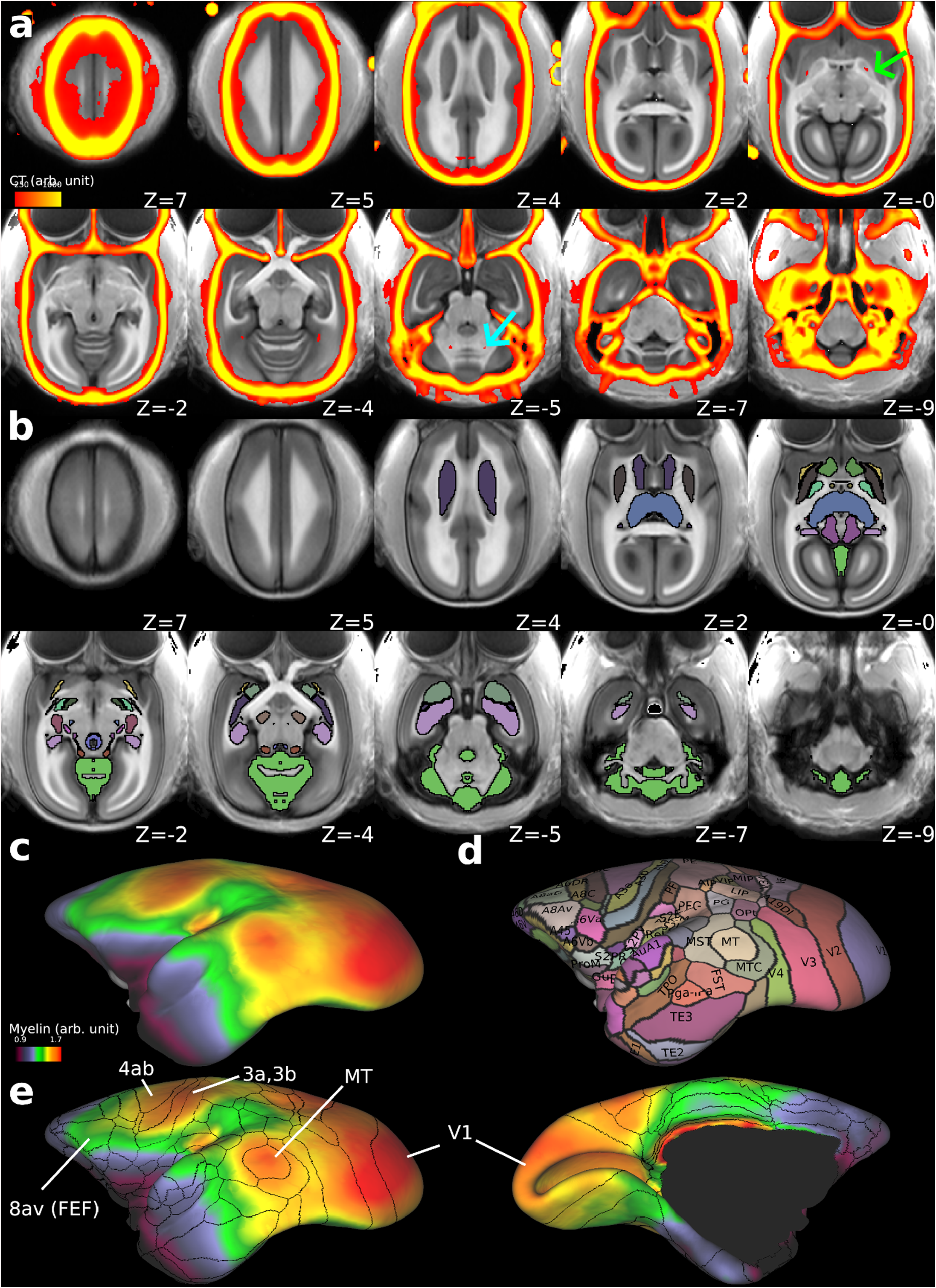
The MarmosetRIKEN20 multi-modal templates (version 1.0) in AC-PC and grayordinates. **(a)** The multi-modal templates with T1w (greyscale, N = 20) overlaid with thresholded CT (red-yellow, N = 10). Each subject’s CT image was registered to T1w in AC-PC native coordinates using the MBFR+BBR, warped to the AC-PC template coordinates using an MRI warp field, and averaged across subjects. Note that physiological calcifications were found bilaterally in the globus pallidus (green arrow, z = 0) and dentate nucleus (cyan arrow, z = −5). **(b)** the subcortical grey matter atlas of MarmosetRIKEN20 including 21 subcortical grey matter regions, and anterior and posterior commissures (colour coded, outlined by black line) overlaid on the T1w template image (grey colour). Readers can find annotations of each region by accessing the data at BALSA database. **(c)** T1w divided by T2w myelin map overlaid on the average midthickness surfaces of MarmosetRIKEN20. **(d)** Surface version of marmoset cortical parcellation atlas of Paxinos, Watson, Petrides, Rosa and Tokuno (Paxinos et al., 2012) including 116 cortical areas. **(e)** Outlines of the cortical parcellations overlaid on the myelin map (left, lateral; right, medial view). Note that high myelin contrast is colocalized with the parcellations at MT, somatomotor sensory areas (4ab, 3a, 3b) and visual cortex (V1), and relatively high myelin with the area 8av, frontal eye field (FEF). Data at BALSA: https://balsa.wustl.edu/study/p005n

### 3.6. Application to image-guided neurosurgery

Distinct neurosurgical strategies guiding surgical instruments for operations into cortical or subcortical locations are illustrated in Figure 9. For subcortical neurosurgery (right), the 3D coordinates (either in template or native coordinates) are selected and the target is warped to the subject’s AC-PC native coordinates. A robot, such as Brainsight, can transform these coordinates online to surgical native stereotactic coordinates. For cortical neurosurgery (left), the target is first identified by the vertex on the cortical midthickness surface in the ‘grayordinate’ system and its 3D coordinates in the subject’s AC-PC native coordinates. Then the robot provides the coordinates in the surgical stereotactic coordinates.

**Figure 9.**
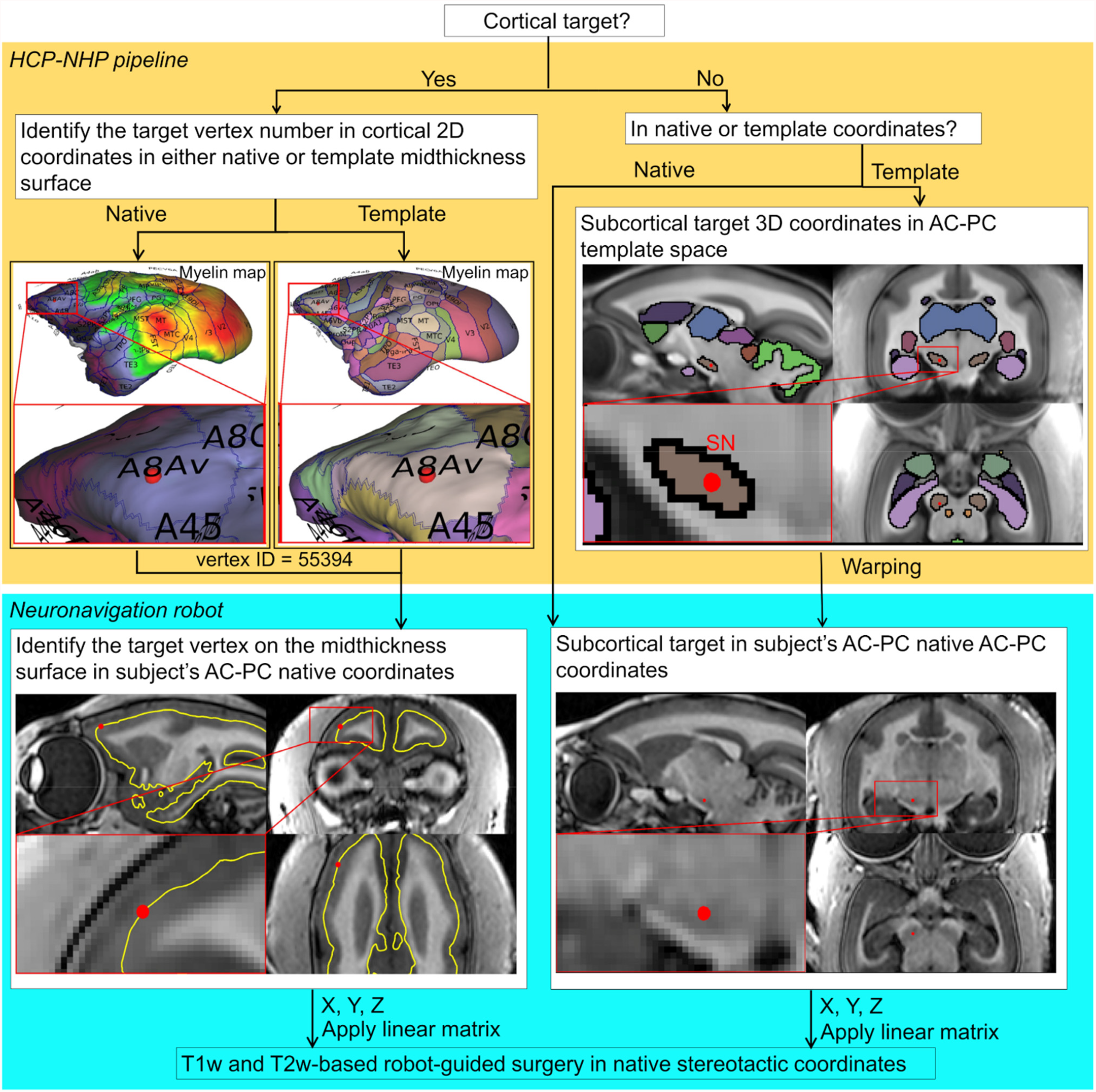
Neuronavigation strategy for cortical and subcortical targets in marmosets. For cortical surgery, the target is first identified on the vertex in cortical 2D coordinates (in either native or template midthickness surface). Next, the 3D coordinates corresponding to the vertex number of interest are read in the subject’s AC-PC native coordinates. For subcortical surgery, the target is identified in the 3D coordinate system (either in template or native coordinates). When the template coordinate system is used, the target’s 3D coordinates are warped to the subject’s AC-PC native coordinates. The robot-guided neurosurgery utilises these 3D coordinates by transforming from subject’s AC-PC native to the stereotactic coordinates. Study ID: (MRI: A17051101)

Comparison between the registration guided surgery plan and postoperative MR image demonstrates precise insertion of the guiding cannula deep into the brain (Fig. 10). The cannula-insertion positions were identified with respect to the native surface of the cranium using Brainsight, and the cannula-insertion trajectories were planned according to the native MRI space to Cau and SN, respectively (Fig. 10b). The coronal sections of the cannula for SN and Cau were tilted in the posterior direction from vertical by 12° and 1°, respectively (Fig. 10c). The postoperative MR image confirmed that the distance between the tip of the guiding cannulas and the targets were 1.6 mm, enabling precise injection cannula insertion for drug delivery (Fig. 10c). The target error in SN was 0.2 mm. However, this experiment was done only in a single subject, and therefore we cannot estimate the consistency with which such precision can be achieved (see Discussion).

**Figure 10.**
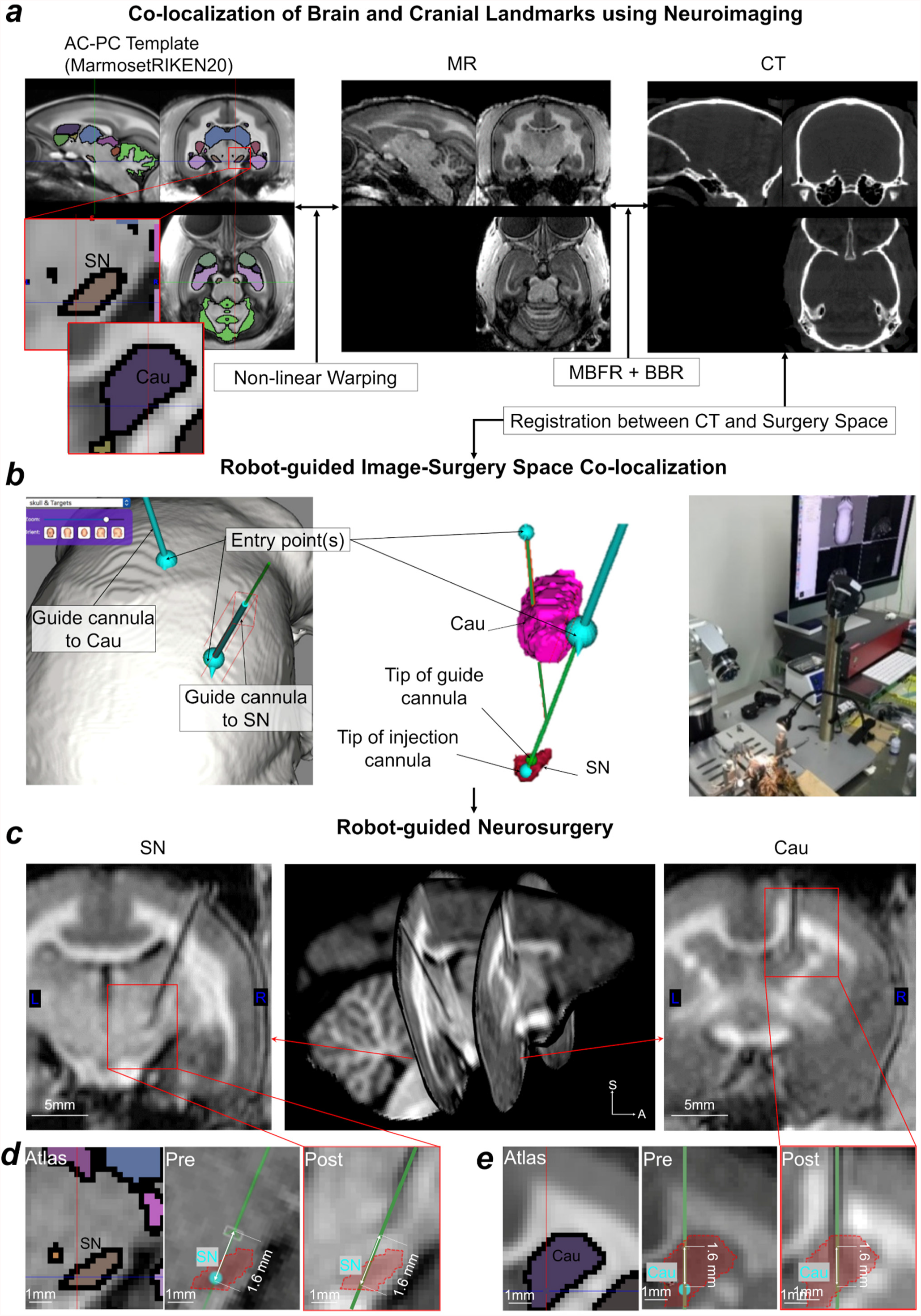
Exemplar application of neuronavigation system. **(a)**The targets were set in the substantia nigra (SN) and the caudate nucleus (Cau) in the AC-PC template coordinates. Template is overlaid with Paxinos atlas (Paxinos et al., 2012).The target locations were non-linearly warped to the subject’s AC-PC native coordinates (left panel to middle panel). The subject’s MR and CT images are pre-registered by the marker-based fiducial registration (MBFR) and then fine-tuned by the boundary-based registration (BBR) methods (middle and right panels). The neuronavigation robot imports the MR or CT images in subjects’ AC-PC native coordinates to navigate to the target in the surgery space. **(b)** The cannula-insertion positions were identified with respect to the location over the surface of the cranium. Presurgical trajectories aiming for Cau (pink) and SN (red) are shown in native MRI space. Note that the cannula-insertion trajectory to SN avoided Cau and ventricles (middle panel). **(c)** Postoperative MR images. Confirmation of guide cannula insertion to SN and to Cau. The tip of the cannula was planned at a position of 1.6 mm from targets, because the needle used for the drug administration extends 1.6 mm from the tip of the cannula. **(d)** Subcortical atlas (in color) registered to AC-PC native coordinates with the target position of the SN (cyan point, middle panel), the planned trajectory in the preoperative MR image (green line, middle panel), and the position of the cannula position (arrow head) and planned trajectory (green line) in the postoperative MRI (right panel). **(e)** Subcortical atlas (in colour) registered to AC-PC native coordinates with the target position of the Cau in AC-PC native coordinates (cyan point, middle panel), the planned trajectory in the preoperative MR image (green line, middle panel), and the position of the cannula position (arrowhead) and planned trajectory (green line) in the postoperative MRI (right panel). Study ID: (CT: 19060401, MRI: A19060401, A19081902)

## 4. Discussion

The marmoset is an increasingly important NHP laboratory model in neuroscience and biomedical research due to its evolutionary proximity to humans relative to intensively studied rodents (Okano et al., 2016), and complex social behaviours (Miller et al., 2016). Recent developments in gene manipulation (Sasaki et al., 2009), functional imaging (Hori et al., 2020; Liu et al., 2019, 2021; Sadakane et al., 2015), white matter pathways and neural tracing (Liu et al., 2020, Majka et al., 2020, 2021), and cellular mapping (Murakami et al., 2018) are also expected to provide evidence how variability of the behaviours are associated with brain and functional segregation and diversity in this species. In this study, we found that common marmosets have substantial intersubject variability in cranial contours and landmarks, size of brain and brain regions, and cortical surface landmarks so that it significantly impacts the choice of coordinate system when experimenters perform brain localization and targeting. Spatial localization of cranial and brain landmarks, commonly used in stereotactic procedures, have substantial uncertainty. This ambiguity arises mainly from anatomical variability in cranial and brain morphology but also from experimental variability in positioning the ear canals and orbital ridges relative to the stereotaxic frame. To overcome these limitations for interventional brain studies, we introduced a methodology utilising marker and boundary-based brain registration and targeting systems in marmosets.

### 4.1 Marker and boundary-based registration

We demonstrated that image alignment using a combination of MBFR initialization and BBR fine-tuning allows more robust and accurate registration as compared to software-only or marker only methods. MBFR was effective for robust initialization, which has been often problematic in software-based registration (Greve and Fischl, 2009; Hill et al., 2001). Indeed, our study showed that software-based registration resulted in frequent initialization failures (Fig. 3). In contrast, MBFR did not result in any failures. The software-based registration methods depend on the accuracy of initialization, whereas the MBFR is only dependent on the accuracy of determination of the centroid of markers. Application of BBR after MBFR further improved the accuracy of registering the cortical surface and bones (Fig. 3b,c). The registration error in MBFR+BBR was larger than MBFR alone which can be attributed to the identification errors of the centroid of the markers depending on the image resolution and asymmetry of the marker and the fact that the MRE itself was measured based on the markers, a form of circularity. The cost function of BBR is reasonably low in all cases, which indicates high reliability and accuracy. Therefore, the MBFR combined with BBR enabled highly accurate registration by compensating for the weak points of each other (the initialization dependency or the marker centroid).

### 4.2 Individual variability of marmosets

Using the multi-modal brain targeting system, we demonstrated substantial intersubject variability of cranial and brain landmarks, particularly the bregma. This intersubject variability causes uncertainty in conventional stereotactic surgery. Bregma location varied by 2.0 mm (in 2SD) in the anterior-posterior direction across marmosets (Table 2, Fig. 6,7a). Previous studies of rodents reported much smaller variability (2SD): ∼0.6 mm in rats (Paxinos and Watson, 2017) and 0.5 mm in mice (Paxinos and Franklin, 2019). Larger intersubject variability of the marmoset bregma may be ascribed to larger cranial size or to larger individual variability than in rodents, but we consider the former is not likely the case. To normalise differences in scales and dimensions, we calculated the isometric ratio of brain scales and variability (Hayashi et al., 2021) (Fig.11). The results disclosed that marmosets had high variability of bregma (4.0) and brain volume (3.6) relative to other rodents (rat 1.2 and 1.6; mouse 1.0 and 1.0 respectively), whereas the size of the brain is only 2-fold difference (marmoset 2.3, rat 1.4, mouse 1.0). Taken together, these findings indicate the intersubject positional variability of bregma and size variability of the brain are approximately 3 to 4-fold larger in marmosets than in laboratory rodents and can be ascribed largely to high intersubject variability rather than large brain scale in primates.

**Figure 11.**
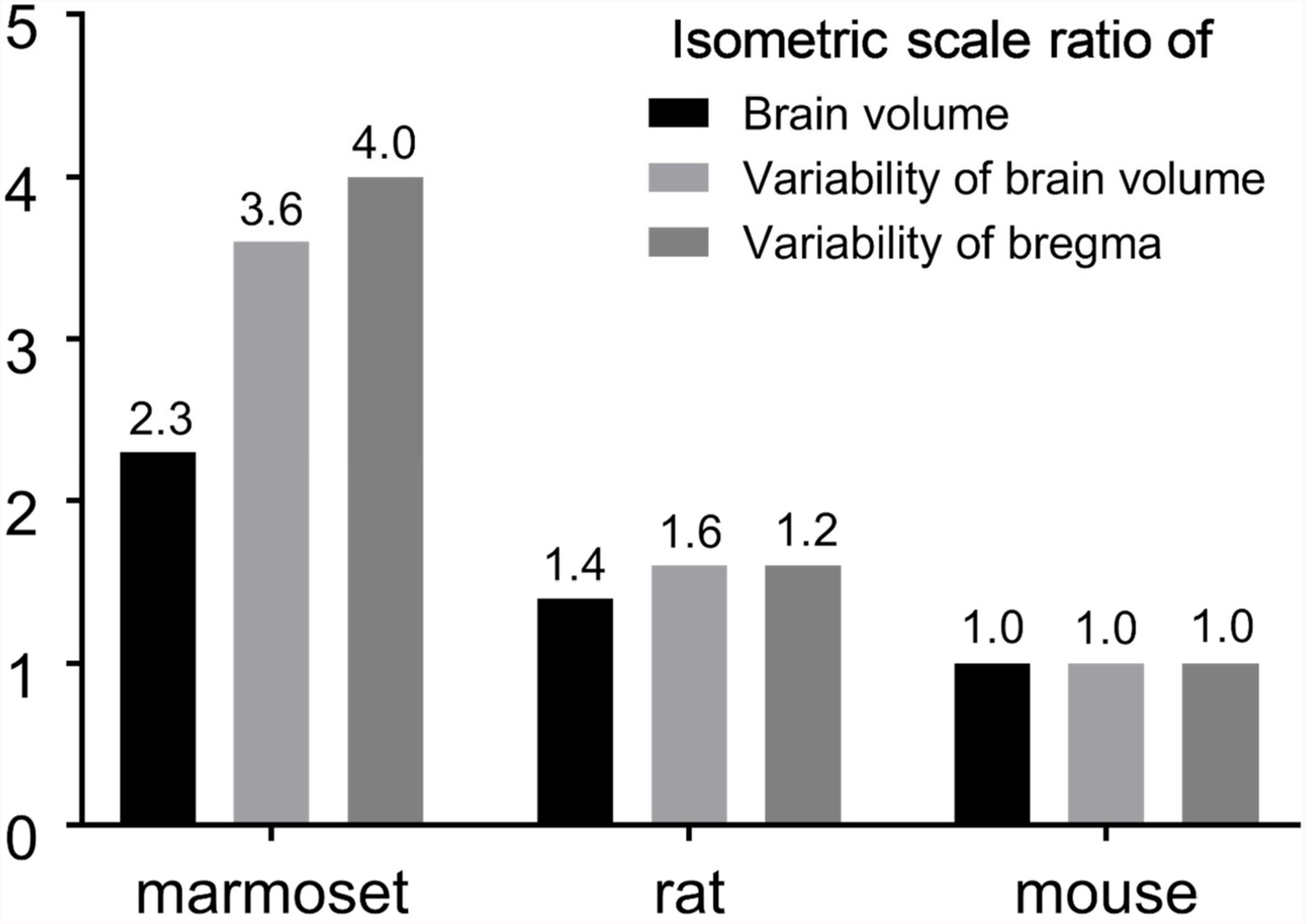
Variability of the bregma and brain volume across species. The variability of the bregma is larger in marmosets compared to rats and mice. Thus, the bregma is not recommended as a cranial landmark in marmoset neurosurgery. The isometric scale ratio of the 3D brain volume (or variability) was calculated using the ratio between the cube root of each species’ brain volume (or its variability) relative to that in mice. The isometric scale ratio of bregma variability in the Y-direction was calculated using the ratio between bregma variability in each species relative to that in mice. All the marmoset data is from the current study, whereas the rodent bregma data from Paxinos and Watson, 2017 and Paxinos and Franklin, 2019, and brain volume data from Hasegawa et al., 2010 and Ma et al., 2008. Note that all the data is from animals housed for experimental use, but sampling is not harmonised (e.g., age and degree of inbreeding) across species.

The intersubject variability of cortical landmarks is also significant in marmosets. The variability of brain organisation is of particular interest in this species as social behaviours and personality are significantly variable (Yokoyama et al., 2013; Miller et al., 2016; Okano et al., 2016; Yokoyama et al., 2021). Since major functions of the inferior parietal region include motor coordination, spatial perception, visual attention, and social perceptions (Sui et al., 2015), the structural and functional variability of the inferior parietal region may also impact subject-specific behavioural capabilities (Mikula et al., 2007; Pomberger et al., 2019; Yokoyama et al., 2021). Our results demonstrated that the location and presence of the marmoset IPS also significantly varies between subjects and hemispheres. Since the asymmetry of IPS is also found in its ramification patterns of the human brain (Zlatkina, V., Petrides, M., 2014), it is of interest to investigate the variability of cortical functional areas and morphology of marmoset in future studies.

Marmoset electrophysiological recordings (Rosa et al., 1997; Rosa and Elston, 1998; Rosa and Schmid, 1995) and tracer injections (Burman et al., 2014, 2006; Reser et al., 2013) have often used cortical landmarks when targeting the needles (e.g., IPS, calcarine sulcus, lateral sulcus, superior temporal sulcus). However, the precise coordinates of these landmarks are not easily defined in their exact spatial location on the cortical sheet. For example, our results indicate that the IPS exists in either or both hemispheres fewer than half of animals investigated (section 3.4). In addition, the location of the deepest point of IPS was variable across subjects in the y direction by over 5% of anterior-posterior length of the average brain size (Table 2), which is comparable with those in humans (variability of location in y direction of the central and superior temporal sulcus was 10-14 % of the brain) (Steinmetz et al., 1990). We also found significant cross-subject variability in the termination of the lateral fissure and in sulcal depth of superior temporal sulcus (data not shown), regions that are involved in social behaviours (Suzuki et al., 2015).

Taken together, we conclude that cranial and brain surface landmarks are not precise reference points to guide stereotactic surgery in the small NHP brain. Instead, we argue that surgical planning in small NHPs may benefit from being specifically designed according to each subject’s own anatomy in a grayordinate system based on multi-modal neuroimaging data. Recent human neuroimaging studies suggest that intersubject alignment based on the highly variable folding/sulcal patterns fail to achieve accurate cortical functional localization (Glasser et al., 2016; Coalson et al., 2018), whereas multi-modal registration using functional connectivity as well as myelin content more accurately compensates for individual variability.

### 4.3 Grayordinates, templates and atlases of marmoset brain

To account for cortical intersubject variability in cortical curvature and total surface area, we introduced a CIFTI grayordinate space (Glasser et al., 2013) for common marmosets. The primary purposes of the grayordinate space are to respect the topology of the cortical sheet, whether lissencephalic or gyrencephalic, to explicitly map cortical data onto a standardised 2D surfaces (Glasser et al., 2013). This approach has already been applied to macaque brains (Autio et al., 2020; Donahue et al., 2018), which enables surface areal feature-based registration and comparison of cortical features across species. The cortical surface approach also enables more precise registration across subjects using the Multi-modal Surface-Matching (MSM) algorithm (Robinson et al., 2018, 2014). This grayordinate-based approach in marmosets may also be advantageous for handling subject variability of cortical folding (e.g., IPS Fig. 7b,c) and functional areas (Glasser et al., 2016).

The multi-modal templates and atlases of the marmoset in grayordinates and volume space are presented. The templates included standardised CT and MR images so that both cranial and brain landmarks are visible in the averaged template space. Very small physiological calcifications in the pallidum and cerebellar nuclei are visible in the CT template, as found in the human brain (de Brouwer et al., 2021), whereas the bregma is not clearly seen due to subject variability (Fig. 6). The cortical areal and subcortical volume parcellations were created based on the image contrasts of T1w and T2w and the histology atlas of Paxinos et al. (2012). We found the close similarity between the myelin contrast (a ratio of T1w divided by T2w) and the the cortical parcellations of some cortical areas (e.g., visual, somatosensory, auditory, MT, and FEF), as well as T1w and T2w contrasts of many subcortical volume structures (e.g., basal ganglia, thalamus, periaqueductal grey, habenular nucleus, lateral and medial geniculate nucleus). However, further validation of brain parcellations are needed in future studies by combining histology data (e.g., Majka et al., 2021), as well as functional connectivity data as has been done in humans (Glasser et al., 2016). This may require refinement of technologies in terms of spatial mapping of 2D-histology data into 3D neuroimaging data (Wang et al., 2020; Majka et al., 2021; Hayashi et al., 2021) and intersubject registration based on multi-modal data for cortical surfaces (Robinson et al., 2018) and brain volume (Lange et al., 2020).

### 4.4 The size of the eyeball and pitch angle between AC-PC and stereotactic coordinates

We found significant bias of brain coordinates between traditional stereotactic and intracerebral landmarks (AC-PC). The marmoset stereotactic horizontal plane is tilted by + 10.0° ± 1.3 pitch (i.e., frontal *downward* direction by 10.0°) relative to the AC-PC plane (Fig. 4). This result conflicts with a prior report that these coordinate systems have the same orientation (Risser et al., 2019). Importantly, this pitch bias is different across species, as the macaque stereotactic plane is tilted by – 3 to – 15° pitch (i.e., frontal *upward by 3 to 15*°) relative to AC-PC plane (Jung et al., 2021; Klink et al., 2020). Moreover, in humans, the stereotactic plane is approximately tilted by −10° pitch (i.e., frontal *upward* direction by 10°) with respect to the AC-PC plane (Park et al., 2010). This coordinate system bias across species may originate from the size of the eyeball relative to that of the brain and other factors such as shape of the cranium and face. For example, species with large eyes relative to the brain size (e.g., marmosets) tend to have a large or positive pitch (frontal *downward* direction) of the stereotactic coordinate relative to AC-PC plane, whereas species (e.g., human and macaque) with relatively smaller eyes tend to have a smaller or negative pitch (frontal *upward* direction). Indeed, the volume ratio of eyeball to brain is substantially larger in marmoset (10%) (Korbmacher et al., 2017) in comparison to macaque monkeys (3%) (Atsumi et al., 2013) and humans (0.4%) (Heymsfield et al., 2016).

### 4.5 Neuroimage-guided neurosurgery

The combination of MBFR and BBR enabled robust and accurate cross-modal registration between MRI, CT, and the surgical device (Figs. 3,9). MBFR provided a 100% success rate and performed well as an initialization for BBR. Thus, the combined approach achieved reproducible and accurate registrations across subjects (Fig. 3). Indeed, this result was supported by the accurate proof-of-concept insertion of cannula into the deep brain structures (Fig. 10). Importantly, the cross-modal registration method can overcome cranial (e.g., bregma location) and brain (e.g., IPS location) subject variability in marmosets (Table 2). Taken together, the combination of MBFR and BBR registration may be a practical as well as accurate tool for image-guided neurosurgery.

High registration accuracy is critical to neurosurgery of small regions/areas. Since the average marmoset cortical hemisphere surface area is 9.9 cm^2^ (Hayashi et al., 2021) and number of cortical areas is ∼116 (Paxino et al., 2012), the average cortical area size is ∼8.5 mm^2^ (minimum of 0.18 mm^2^ in area 25). This finding suggests that the minimum desired accuracy to target the smallest cortical areas is 0.4 × 0.4 mm. Our proof-of-concept MBFR and BBR-guided robot microsurgery suggests that such accuracy may be achieved even for deep brain structures (error ∼0.2 mm, Fig. 10). For perspective, the average macaque monkey cortical hemisphere area is 106 cm^2^ (Hayashi et al., 2021), number of estimated cortical areas is ∼130-140 (Van Essen et al., 2012) and the average cortical area size is ∼70 mm^2^ (minimum of 5 mm^2^) (Autio et al., 2020). Since frameless neuronavigation system surgery error is 1.05 to 1.2 mm in macaques (Frey et al., 2004; Sudhakar et al., 2019; Zhu et al., 2019), surgical accuracy of the targeting relative to the average cortical area may be comparable in macaque and marmoset (error / average cortical area ≈ 2%). However, it should also be noted that the requirement for final targeting precision depends on the size of the target, which may differ across applications (e.g., electrophysiological recording, microelectrode stimulation, or infusion of drugs, tracers, viral vectors). For example, injections of solution are likely dependent on size of the injected reagent: for viral vectors, volume spread is the same as or a bit larger (1.5 times) than the volume of injected solution (Watakabe et al., 2015), whereas it is much larger (20-30 times) when used with small-molecule drugs like muscimol (Murata et al., 2015). In electrophysiology, the spatial size of a single neuron recording is likely less than 100μm, where that of local field potential (LFP) is reported to be as large as 0.5 to 3mm (Logothetis et al., 2003).

Overall, the image-guided robot microsurgery improved the utility, flexibility and accuracy in a variety of interventional experimental procedures. A similar MRI-guided approach was recently demonstrated in the marmoset (Mundinano et al., 2016), however, this approach required that the surgery was immediately performed after the MRI scan to ensure the same position of the stereotactic device during the procedure. In addition, the stereotactic injection can only be performed in a limited range of angles restricting operational degree of freedom. The operational range is an important factor, for example, when the trajectory of the cannula needs to circumvent lateral ventricle (as in the case of targeting SN, Fig. 10). Since a cannula penetrating a ventricle may increase the target error by multiple passes through the brain (Zrinzo et al., 2009), here we planned the operation by avoiding ventricles at an optimal angle and demonstrated successful targeting of the SN. Future studies should assess the effect of image-guided microsurgery by a larger number of cases and by histological verification of injection sites. Finer interventional techniques (e.g., ultra-fine cannula and needle) may also need to be developed for more accurate targeting. Even if the experimenters have no direct access to MR and CT, the multi-modal templates may be useful for more accurate targeting by taking into account the bias between stereotactic and AC-PC spaces.

## 5. Conclusion

In this study, we evaluated the accuracy of different brain coordinate systems to target specific brain structures in marmosets. We found substantial intersubject variability in brain and cranial landmarks in marmosets. In particular, the variabilities of the cranial landmarks (e.g., bregma, interauricular line) are substantial enough to bias the brain orientation in surgical interventions. Thus, it is recommended to use the brain image and/or cranial landmarks for spatial localization. The population-based volume and surface templates and atlas in grayordinates were created for the first time in marmoset monkeys, which may provide a basis for accurately combining function and histology data in future.

## Abbreviations

AC-PC: Anterior commissure-posterior commissure
BBR: Boundary-based registration
Cau: Caudate nucleus
CIFTI: Connectivity InFormatics Technology Initiative
COV: Coefficient of variation
CSF: Cerebrospinal fluid
CT: Computed Tomography
FEF: Frontal eye field
FLIRT: FMRIB’s Linear Image Registration Tool
FMRIB: Functional Magnetic Resonance Imaging of the Brain
FNIRT: FMRIB’s Nonlinear Image Registration Tool
FOV: Field-of-view
GIFTI: Geometry format under the Neuroimaging Informatics Technology Initiative
HCP: Human Connectome Project
IPS: Intraparietal sulcus
MarmosetRIKEN20: RIKEN marmoset MRI & CT template
MBFR: Marker-based fiducial registration
MRE: Marker registration error
MRI: Magnetic Resonance Imaging
MSM: Multi-modal Surface-Matching
MT: Middle temporal area
NHP: Non-human primate
HCP-NHP: human connectome project non-human primate
NMI: Normalized mutual information
SN: Substantia nigra
T1w: T1-weighted MRI
T2w: T2-weighted MRI
V1: primary visual cortex

## Notes

Supplementary Information is available in the online version of the paper.

## Credit authorship contribution statement

Conceptualization: HW, TH; Methodology: TO, MFG, TH; Software: TO, AU, JAA, HW, TH; Formal analysis: TO, JAA, TH; Investigation: TO, MO, AK, CT, YH, KN, TN, CY; Writing -original draft: TO, JAA, TH; Writing -review & editing: JAA, AU,DVCE, MFG, TH; Visualisation: TO; Resources: HN, TY, CY; Supervision: HW, TH; Project administration: TH; Funding acquisition: TO, TH, DVCE.

## Acknowledgments

We thank Akiya Watakabe, Misako Komatsu, Takuro Ikeda, Takashi Azuma, Yuki Matsumoto, Kenji Mitsui, Reiko Kobayashi, Hanako Hirose and Masataka Yamaguchi for their technical support. This research is supported by the program for Brain/MINDS-beyond (JP21dm0307006, T.H.) and Brain/MINDS (JP21dm0207001, T. Y.) from Japan Agency for Medical Research and development, AMED, JSPS KAKENHI Grant Number (JP15K08707, T.O., JP20K15945, J.A.A. JP18K06372, C.Y.) and NIH Grant MH060974 (D.C.V.E.).

