## Supplementary Fig for "Anatomical variability, multi-modal coordinate systems, and precision targeting in the marmoset brain"

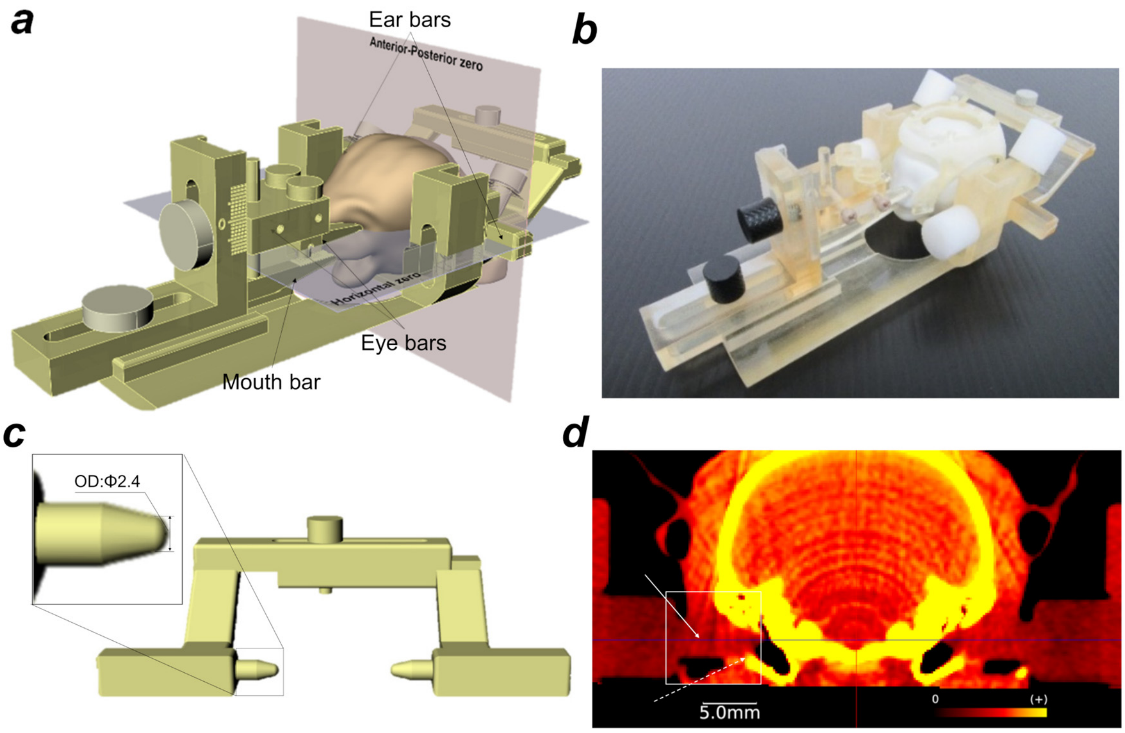


**Supplemental Figure 1.** Design of stereotactic device (a) The device was designed to be compatible with small animal CT (FOV = diameter 73 mm × height 57 mm). The head of the marmoset is mounted on the stereotactic device using mouth, eye and ear bars. (b) The marmoset phantom and head holder are firmly attached to the stereotactic device. (c, d) The tip of the ear bar matched the marmoset's auditory canal. (d) Coronal CT image. Solid and dashed arrows indicate the ear bars and the external auditory canals, respectively.

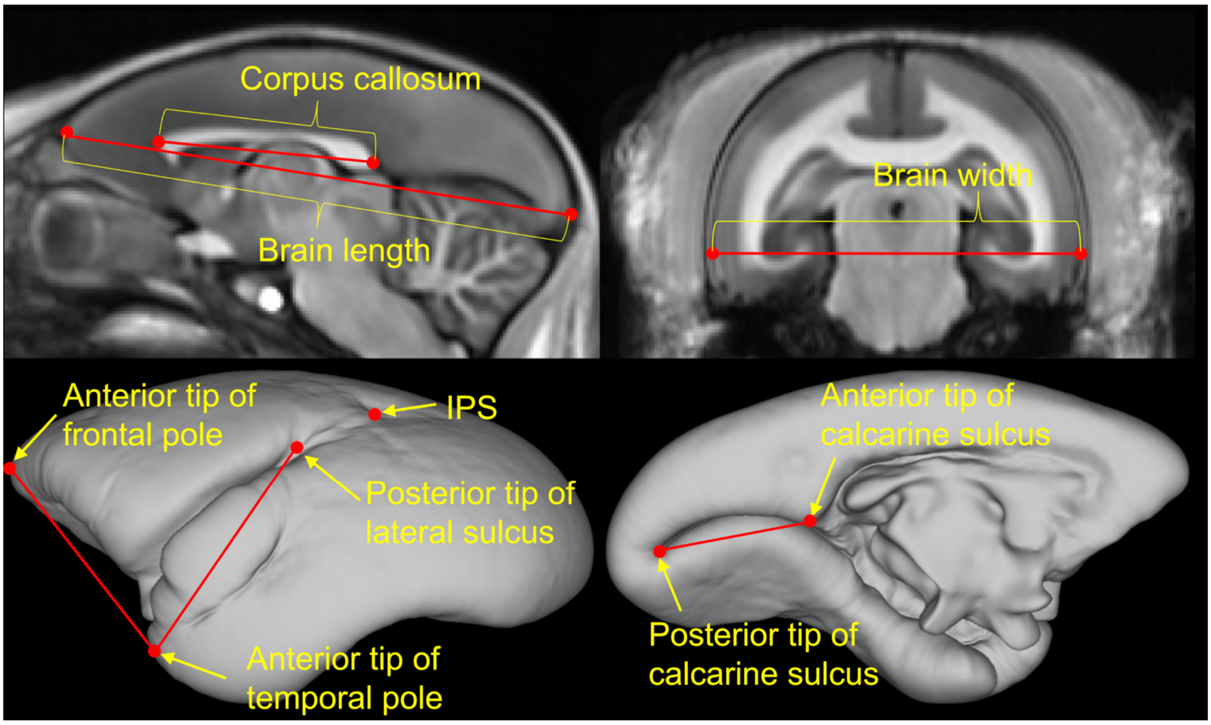


**Supplementary Figure 2**
The figure shows a schema of landmarks and distances of interest. The coordinates of landmarks were identified in T1w AC-PC native space and the Euclidean distances were measured in each animal.
